# Decorin Promotes Nascent Proteoglycan Retention in Cartilage Matrix by Strengthening Collagen II-Aggrecan Integration

**DOI:** 10.1101/2025.11.12.688145

**Authors:** Thomas Li, Gabriela Canziani, Yuchen Liu, Neil Patel, Biao Han, Mingyue Fan, Annie Porter, Bryan Kwok, Chao Wang, Qing Li, David E. Birk, Renato V. Iozzo, Irwin M. Chaiken, X. Lucas Lu, Robert L. Mauck, Lin Han

**Author notes:** Correspondence and requests for materials should be addressed to: Dr. Lin Han.

## Abstract

Cartilage extracellular matrix (ECM), a hydrated collagen II-aggrecan composite, undergoes dynamic turnover during both normal homeostasis and disease-associated remodeling. This study elucidates a crucial role for decorin in promoting the retention and stability of nascent aggrecan within this matrix. By applying bio-orthogonal click-labeling, we demonstrate that loss of decorin accelerates the release of nascent aggrecan under both physiological and inflammatory conditions, without affecting its preferred localization to the pericellular matrix. Conversely, supplementation with exogenous decorin mitigates inflammation-induced loss of nascent aggrecan, supporting its potential as a therapeutic target. At the molecular level, decorin exhibits strong binding affinity for aggrecan, and enhances aggrecan-aggrecan and aggrecan-collagen II interactions, reinforcing its direct role in integrating cartilage matrix constituents. Also, by binding to collagen II, decorin stiffens the collagen II fibril network, thereby strengthening the confinement effect that limits the diffusive loss of entrapped aggrecan. Notably, decorin does not alter chondrocyte transcriptomic profiles *in vivo*, emphasizing its primary role in maintaining matrix integrity through biophysical mechanisms rather than cell signaling. Together, these findings provide a mechanistic foundation for developing decorin-based biomaterials or gene therapies aimed at preserving or regenerating the cartilage matrix for improved outcomes in osteoarthritis.

## 1. Introduction

Osteoarthritis (OA), the most prevalent musculoskeletal disease, afflicts 595 million of the population worldwide,^[1]^ and is hallmarked by the irreversible breakdown of articular cartilage.^[2]^ Successful regeneration of cartilage requires the restoration of its extracellular matrix (ECM), a hydrated composite primarily composed of type II collagen fibrillar network and the large proteoglycan aggrecan.^[3]^ Proper cartilage biomechanical function and chondrocyte mechanotransduction depend on appropriate assembly and integration of these two constituents.^[4–5]^ Within the ECM, aggrecan forms aggregates with hyaluronan (HA)^[6]^ via link protein-assisted binding.^[7]^ These aggregates are entrapped within the collagen fibrillar network in a highly compressed conformation, resulting in a ≈ 50% molecular strain in unloaded tissue.^[8]^ In turn, aggrecan and its chondroitin sulfate-glycosaminoglycan side chains (CS-GAGs) provide the ECM with a negatively charged osmotic environment with low hydraulic permeability, which governs tissue compressive and poroelastic energy dissipative properties.^[9–10]^ Additionally, the collagen II fibrillar network and aggrecan synergistically contribute to the nonlinear tissue tensile and shear properties^[11]^ via a rigidity percolation framework.^[12]^

Cartilage ECM is a dynamically adaptive composite governed by the synergistic activities of cells, matrix molecules and external stimuli. During normal homeostasis, aggrecan undergoes much more rapid turnover than collagen II,^[13]^ with newly synthesized aggrecan predominantly localized in the pericellular matrix (PCM).^[14]^ In human cartilage, aggrecan has an approximate half-life of 4 weeks,^[15]^ compared to about 117 years for collagen II.^[16]^ Despite their vastly different turnover rates, the ECM maintains its integrity throughout lifespan. In early OA, elevated catabolism leads first to aggrecan fragmentation,^[17–18]^ followed by release of aggrecan fragments and degradation of collagen II fibrils.^[19]^ These changes compromise matrix integrity,^[20]^ disrupt chondrocyte mechanotransduction^[21]^ and ultimately result in irreversible cartilage breakdown.^[22]^ However, our understanding of the molecular events that regulate the dynamic turnover and disease-associated degenerative changes of cartilage ECM remains limited, representing a key barrier to developing effective cartilage repair and OA intervention strategies.^[23]^

Our recent work identified decorin, a class I small leucine-rich proteoglycan (SLRP), as a potential regulator of this process. In cartilage, decorin is present at a molar concentration (≈ 15 nmol/mL) comparable to that of aggrecan (≈ 20 nmol/mL),^[24]^ and is actively expressed by chondrocytes throughout human lifespan.^[25–26]^ In early OA, decorin is significantly upregulated^[27–28]^ but not increasingly released from degrading cartilage.^[29]^ This indicates a potentially active role for decorin in ECM remodeling under both normal and pathological conditions. In support, we previously showed that in a young adult decorin-null (*Dcn^−/−^*) murine model, articular cartilage developed reduced aggrecan content (Fig. 1a), impaired load-bearing and energy dissipation properties,^[30]^ and in turn, perturbed chondrocyte *in situ* mechanosensing.^[31]^ Following the destabilization of the medial meniscus (DMM) surgery,^[32]^ decorin ablation resulted in accelerated aggrecan loss, surface fibrillation and more severe cartilage damage,^[33–34]^ highlighting its protective role in OA. Despite these findings, we have not yet uncovered how decorin modulates the dynamic assembly and interplay of cartilage ECM constituents. Such knowledge may provide new strategies for enhancing cartilage repair and preventing degeneration through molecular modulation of decorin activities.

**Figure 1.**
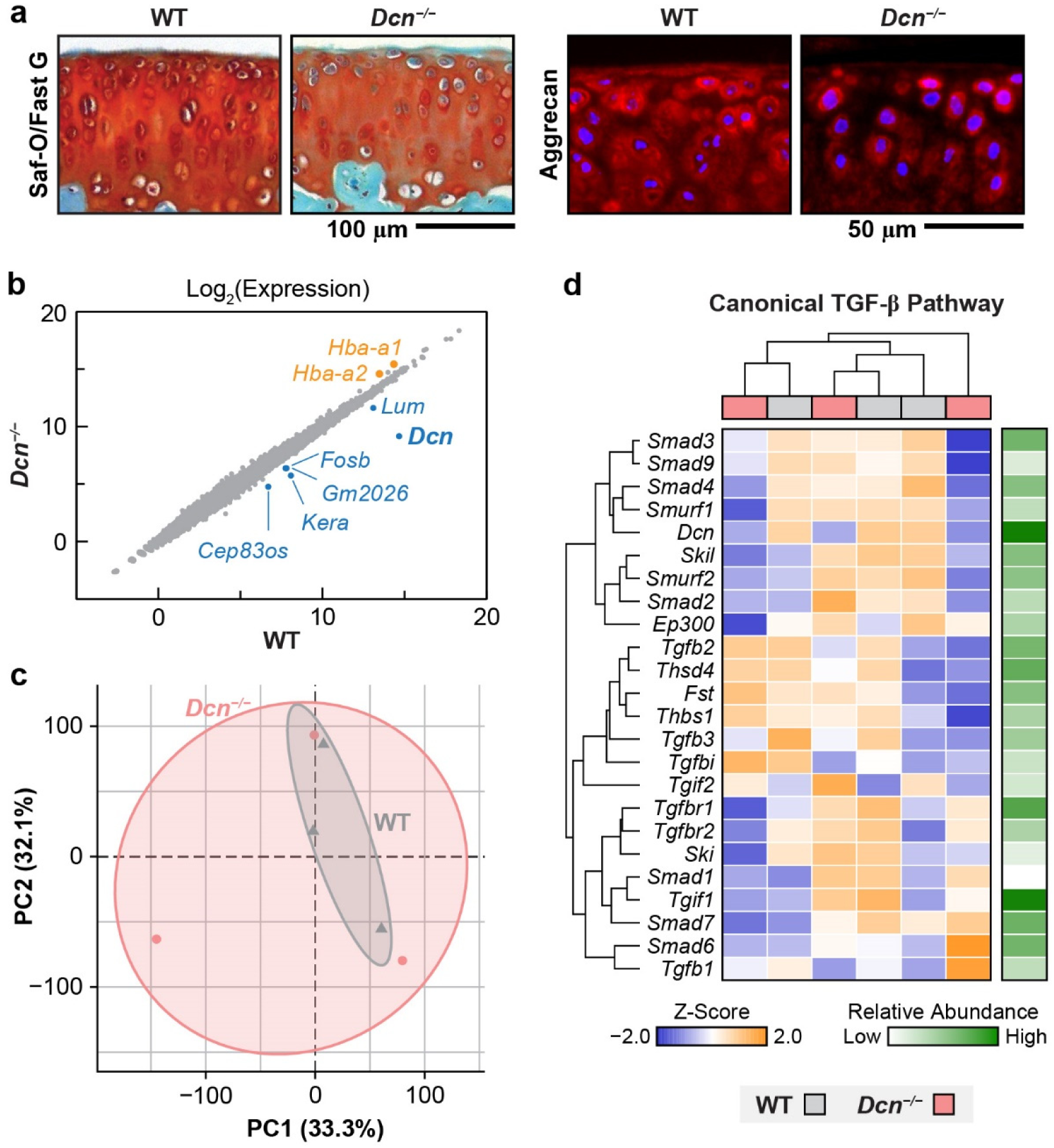
Bulk RNA sequencing analysis reveals no major changes in signaling pathways of decorin-null (*Dcn^−/−^*) cartilage. a) Safranin-O/Fast Green and aggrecan immunofluorescence (IF) images showing pronounced reduction of aggrecan and its sulfated glycosaminoglycans (sGAGs) in young adult *Dcn^−/−^* compared to age-matched wild-type (WT) control (adapted from Ref. ^[30]^ with permission). b) Scatterplot of the log_2_-transformed gene expressions between 3-month-old WT and *Dcn^−/−^*femoral condylar cartilage, with significantly upregulated and downregulated genes in *Dcn^−/−^* cartilage highlighted. b) Principal component analysis (PCA) shows substantial overlap in the clustering of WT and *Dcn^−/−^*samples, indicating no global transcriptomic divergence. c) Heatmap of unbiased gene clustering for the canonical TGF-β signaling pathway shows no distinct separation in TGF-β signaling activities between WT and *Dcn^−/−^*samples (*n* = 3 per genotype).

This study aims to elucidate the molecular mechanisms by which decorin regulates the dynamic turnover and integrity of cartilage ECM and to assess its potential as a therapeutic target for preserving degenerative cartilage. First, given that aggrecan is markedly reduced in *Dcn^−/−^*cartilage (Fig. 1a) and that decorin engages a broad interactome including matrix molecules such as collagens,^[35]^ various growth factors, cytokines and cell surface receptors,^[36]^ we assessed the potential biological effects of decorin on chondrocytes. We performed RNA sequencing to determine how loss of decorin influences chondrocyte transcriptomics and major signaling pathways *in vivo*, such as transforming growth factor (TGF)-β.^[37]^ Next, we applied the emerging bio-orthogonal “click-labeling” technique^[38–40]^ to track the biosynthesis, distribution and retention of newly synthesized proteoglycans and proteins. Studying cartilage explants of *Dcn^−/−^*mice,^[41]^ we investigated the impact of decorin loss on matrix turnover under both normal and degenerative conditions, as well as the potential rescue effects of exogenous decorin in this process. Then, we delineated the molecular interaction kinetics among decorin, aggrecan and collagen II, and queried whether decorin facilitates the integration of aggrecan and collagen II fibrils using surface plasmon resonance (SPR) and atomic force microscopy (AFM)-based nanoindentation. Collectively, our findings demonstrate that decorin plays an indispensable role in cartilage matrix turnover and maintenance by mediating matrix biophysical interactions that strengthen collagen II-aggrecan integration.

## 2. Results

### 2.1. Loss of decorin does not directly alter chondrocyte signaling in young adult cartilage

To test whether the reduction of aggrecan content in *Dcn^−/−^*cartilage (Fig. 1a) stems from decorin influences on chondrocyte signaling and biosynthesis *in vivo*, we applied bulk RNA-sequencing to compare the transcriptomic profiles of chondrocytes isolated from 3-month-old young adult WT and *Dcn^−/−^* cartilage. By this age, *Dcn^−/−^* cartilage has already developed pronounced matrix phenotypic changes, including reduced GAG content, impaired biomechanical properties and altered chondrocyte intracellular calcium signaling *in situ*.^[30–31]^ Among more than 15,000 genes analyzed, we found only eight differentially expressed genes (DEGs) with adjusted *p*-values < 0.05 (Fig. 1b). Also, principal component analysis (PCA) revealed no distinct clustering between WT and *Dcn^−/−^* groups (Fig. 1c), indicating that chondrocytes from both genotypes exhibited largely similar transcriptomic profiles *in vivo*. Notably, although prior studies suggested that decorin may regulate TGF-β signaling by binding to TGF-β,^[37,42]^ we found no evidence of altered canonical TGF-β signaling pathways in *Dcn^−/−^*cartilage (Fig. 1d). Likewise, no differences were observed in other pathways potentially influenced by decorin, including non-canonical TGF-β (Fig. S1),^[43]^ epidermal growth factor receptor (EGFR)^[44]^ and vascular endothelial growth factor (VEGF),^[45]^ Met-mediated hepatocyte growth factor receptor (HGFR) signaling^[46]^ (Fig. S2). Furthermore, gene ontology (GO) analysis did not detect marked genotype-associated differences in matrisome genes (Fig. S3), suggesting that loss of decorin does not directly affect chondrocyte biosynthesis of cartilage matrix content *in vivo*. These results were consistent with our previous work showing *Dcn^−/−^* chondrocytes did not exhibit altered biosynthesis of aggrecan compared to the WT in response to TGF-β stimulation in 3D culture *in vitro*.^[30]^ Therefore, these results indicate that the primary role of decorin in cartilage is to maintain matrix integrity, rather than to directly regulate chondrocyte signaling in young adult tissue.

### 2.2. *Dcn^−/−^* explants exhibit reduced retention of nascent GAGs, but not nascent proteins

To query the role of decorin in dynamic turnover and maintenance of cartilage matrix, we assessed the effects of decorin loss on the distribution and retention of newly synthesized proteoglycans and proteins in an explant model. Using femoral head cartilage explants from 3-week-old wild-type (WT) and *Dcn^−/−^*mice, we applied bio-orthogonal metabolic labeling for 3 days following the 2-day pre-culture immediately after harvest (Fig. 2a). The azide-modified monosaccharide *N*-azidoacetylgalactosamine-tetraacylated (GAL) was used as an analog of *N*-acetylgalactosamine (GalNAc) to label newly synthesized glycosaminoglycans (GAGs), and azidohomoalanine (AHA), a methionine analog, was used to label newly synthesized proteins (Fig. S4).^[40]^ Given that GalNAc is primarily incorporated into chondroitin sulfate (CS)-, dermatan sulfate (DS)-, and to a lesser extent, heparan sulfate (HS)-GAGs,^[47]^ and the majority of GAGs in cartilage are CS-GAGs on aggrecan,^[3]^ GAL signals mainly represent the CS-GAGs of nascent aggrecan. Similarly, since collagen II is the major protein of cartilage,^[3]^ AHA signals primarily reflect newly synthesized collagen II. Subsequently, we fluorescently tagged nascent GAGs or proteins from explants separately labeled by GAL and AHA via bio-orthogonal click-labeling with dibenzocyclooctyne (DBCO),^[40]^ and then cultured the explants with or without 10 ng/mL of the inflammatory cytokine IL-1β for six days (Fig. 2a).

**Figure 2.**
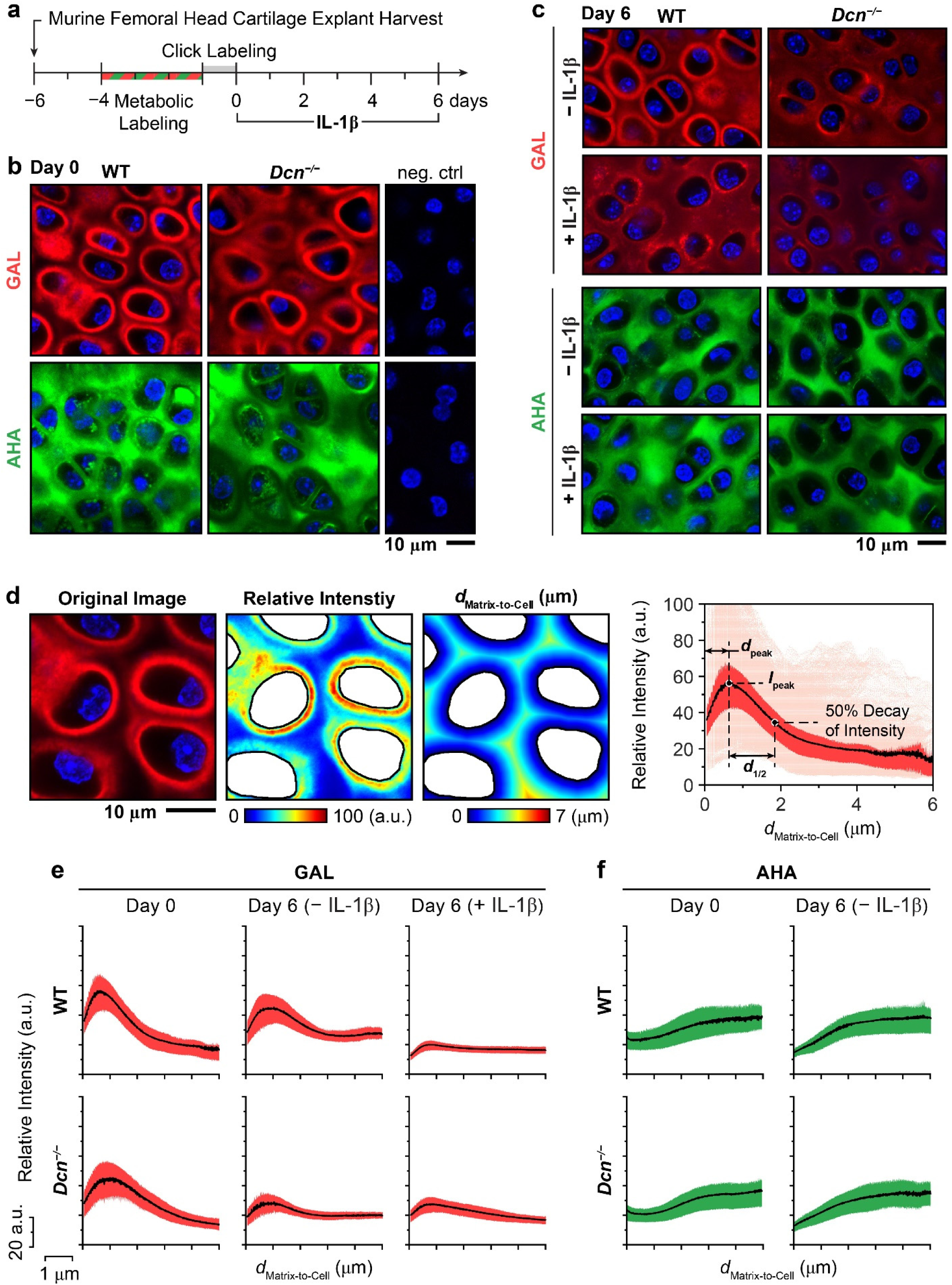
Spatial distribution of newly synthesized glycosaminoglycans (GAGs) and proteins in native cartilage matrix via bio-orthogonal click-labeling. a) Schematic of the experimental timeline for click-labeling and interleukin-1β (IL-1β) treatment of murine femoral head cartilage explants. b) Representative confocal images of GAL- (red) and AHA- (green) labeling in 3-week-old wild-type (WT) and decorin-null (*Dcn^−/−^*) cartilage explants at day 0 of culture, showing the preferential localization of nascent GAGs in the pericellular matrix (PCM) and broad distribution of nascent proteins throughout the extracellular matrix (ECM). Negative control: explants without metabolic labeling but with DBCO click-labeling (blue: DAPI). c) Representative GAL and AHA confocal images at day 6 of culture show reduced nascent GAGs with IL-1β stimulation in both WT and *Dcn^−/−^* cartilage, and no appreciable changes in nascent protein distribution. d) Left panel: Representative 30 × 30 μm^2^ high-resolution (2,048 × 2,048) confocal image of GAL-labeled WT explant, illustrating the extraction of pixel intensity and corresponding matrix-to-cell Euclidean distance, *d*_Matrix-to-Cell_, maps. Right panel: Corresponding GAL-fluorescence intensity profiles for individual pixels as a function of *d*_Matrix-to-Cell_ within a single ROI, illustrating the definition of spatial parameters, the peak intensity, *I*_peak_, the *d*_Matrix-to-Cell_ corresponding to the peak intensity, *d*_peak_ and the distance from *d*_peak_ to the mid-point between peak and baseline intensities, *d*_1/2_ (as defined in Table 1). Shown are raw data, interquartile range (IQR) and mean values for *n* ≥ 9 cells within the ROI. e,f) Representative spatial distributions of e) GAL and f) AHA signals within each ROI for WT and *Dcn^−/−^* explants immediately after click-labeling (day 0) and after 6 days of culture with or without IL-1β (mean with IQR). Images illustrate distinct spatial distributions of nascent GAGs and proteins, and the accelerated release of nascent GAGs due to decorin loss and IL-1β treatment.

**Table 1.** Glossary of quantitative parameters describing the spatial distribution of nascent glycosaminoglycans (GAGs) and proteins in cartilage

| Parameters | Unit | Definition |
| --- | --- | --- |
| $I_{\text{mean}}$ | a.u. | Mean intensity of the radial fluorescence profile, averaged across all matrix-to-cell Euclidean distances within each region of interest (ROI). |
| $I_{\text{peak}}$ | a.u. | Maximum intensity of the radial fluorescence profile, averaged across all matrix-to-cell Euclidean distances within each ROI. |
| $d_{\text{peak}}$ | $\mu\text{m}$ | Distance from the cell surface at which the fluorescence intensity reaches its maximum. |
| $d_{1/2}$ | $\mu\text{m}$ | Distance from $d_{\text{peak}}$ to the point where the intensity decays by 50%, calculated as the midpoint between the peak and baseline (minimum) intensity. |
| $d_{\text{peak}} + d_{1/2}$ | $\mu\text{m}$ | Distance from the cell surface to the point where the intensity decays by 50% based on the midpoint between the peak and baseline intensities. |

For both genotypes, immediately after click-labeling, nascent GAGs were primarily localized to the pericellular domain, whereas nascent proteins were distributed throughout the intercellular matrix space (Fig. 2b). From day 2 to day 6 of culture, GAL signals for nascent GAGs, particularly in the PCM, progressively diminished, with further reduction observed under IL-1β treatment (Fig. 2c and Fig. S5a). Compared to WT, *Dcn^−/−^* explants exhibited a more pronounced reduction in GAL signals, illustrating accelerated loss of nascent GAGs. In contrast, AHA signals showed no appreciable differences between genotypes or treatment conditions (Fig. 2c and Fig. S5b), suggesting that the distribution and localization of nascent proteins were not affected by decorin loss or IL-1β during the six-day culture duration. Additionally, loss of decorin did not alter the distribution patterns of either nascent GAGs or proteins. These results suggested that decorin was not required for the preferential localization of nascent aggrecan in the PCM, or the widespread presence of collagen II throughout the intercellular space.

We then applied custom-built MATLAB imaging analysis programs to quantify the fluorescent signals of nascent GAGs and proteins, as well as their spatial distributions relative to the matrix-to-cell Euclidean distance within ≈ 100 × 100 µm^2^ regions of interest (ROIs) (Fig. 2d). For GAL-labeling, given the presence of a signal peak in the PCM within each ROI, we calculated the average and peak fluorescence intensity, *I*_mean_ and *I*_peak_, the Euclidean distance corresponding to *I*_peak_, *d*_peak_, as well as the distance from *d*_peak_ to the point the intensity decays by 50%, *d*_1/2_ (Table 1 and Fig. 2d,e). For AHA-labeling, due to the absence of clear signal peaks (Fig. 2f), only *I*_mean_ was quantified per ROI. At day 0, immediately after click-labeling, no significant differences were observed between WT and *Dcn^−/−^*explants in any of the spatial parameters for nascent GAGs, or in *I*_mean_ for nascent proteins (Fig. 3a-d and Fig. S6). For example, *d*_peak_ was 910 ± 60 nm for WT and 915 ± 70 nm for *Dcn^−/−^* explants, respectively, and *d*_1/2_ was 924 ± 81 nm for WT and 817 ± 101 nm for *Dcn^−/−^* explants, respectively.

**Figure 3.**
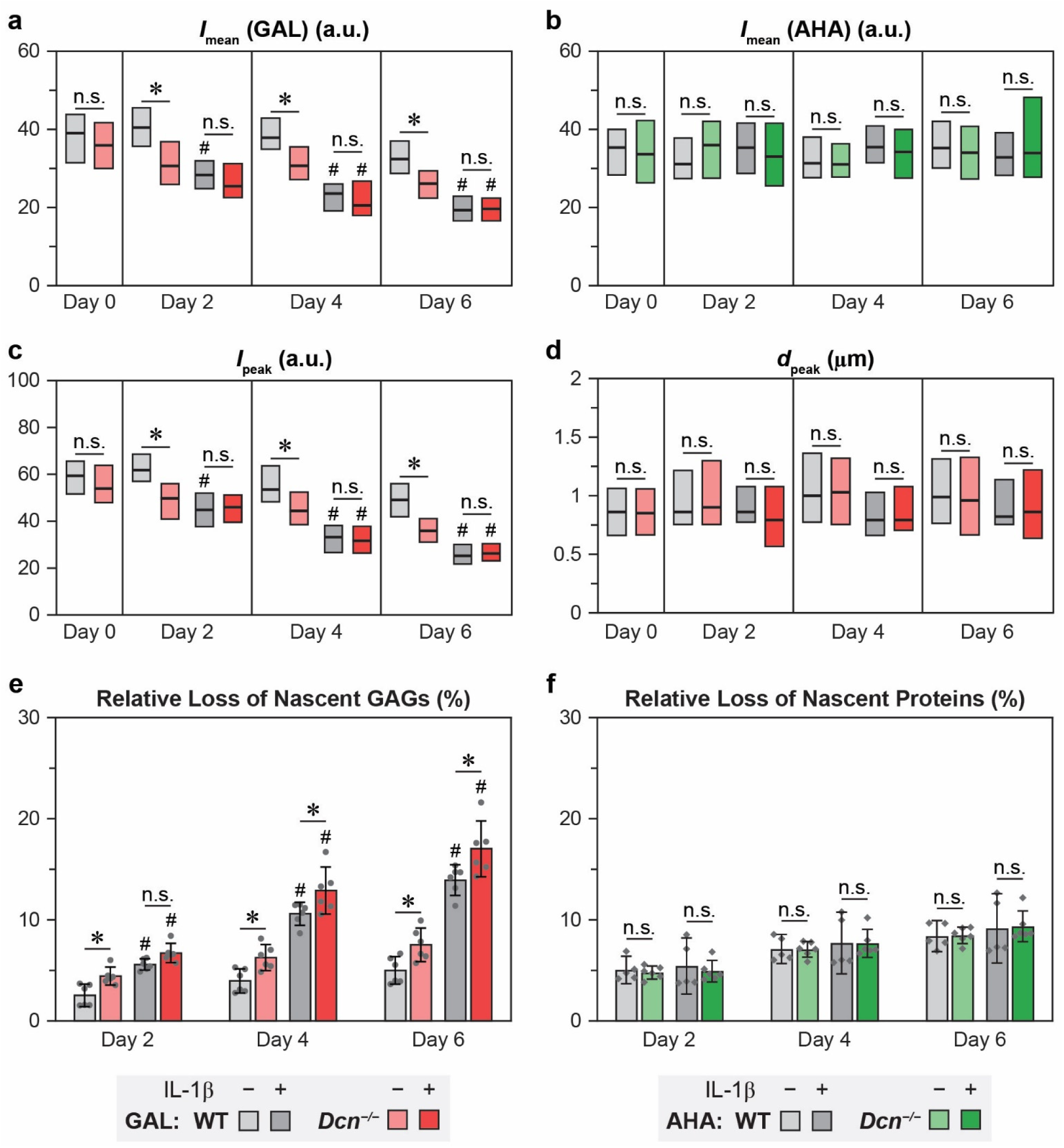
Loss of decorin decreases the retention of nascent GAGs, but not nascent proteins, in cartilage explants with or without the stimulation of IL-1β. a-d) Comparison of spatial parameters for GAL and AHA signals between wild-type (WT) and decorin-null (*Dcn^−/−^*) cartilage from day 0 to day 6 of culture with or without IL-1β: a,b) *I*_mean_ for GAL and AHA signals, respectively, c) *I*_peak_ and d) *d*_peak_ for GAL signals. Box plots for data obtained from > 75 ROIs across *n* ≥ 4 animals per group. e,f) Cumulative release of e) nascent GAGs and f) nascent proteins from WT and *Dcn^−/−^*cartilage explants over 6 days of culture with or without IL-1β. Each data point represents one biological replicate (mean ± 95% CI, *n* = 6). Panels a) to f): *: *p* < 0.05 between genotypes at the same time point, ^#^: *p* < 0.05 versus untreated samples of the same genotype and time point, n.s.: not significant. A complete list of statistical analysis outcomes for *I*_mean_(GAL), *I*_mean_(AHA), *I*_peak_, *d*_peak_, cumulative release of nascent GAGs and proteins is summarized in Tables S1–S4, S7 and S8, respectively.

Over the six-day culture period, *I*_mean_ and *I*_peak_ progressively decreased in both WT and *Dcn^−/−^* explants, evidencing the loss of nascent GAGs. In the absence of IL-1β, *Dcn^−/−^* explants showed accelerated reductions in both parameters vis-à-vis WT, resulting in lower *I*_mean_ and *I*_peak_ for *Dcn^−/−^* explants from day 2 to day 6 (Fig. 3a,c). With IL-1β treatment, both genotypes experienced substantial reductions in *I*_mean_ and *I*_peak_, and as a result, these attenuated signals did not show apparent genotype-associated differences (Fig. 3a,c). Throughout the culture, we found no significant genotype- or treatment-associated differences in *d*_peak_ (Fig. 3d), *d*_1/2_ or *d*_1/2_ + *d*_peak_ (Fig. S6), suggesting that loss of decorin or IL-1β-induced catabolism did not alter the preferred localization of nascent aggrecan or GAGs in the PCM.

We next quantified the relative proportions of nascent GAGs and proteins released into the media as compared to those retained in the explants. Both genotypes exhibited a progressive release of nascent GAGs into the media, with significantly higher release in response to IL-1β stimulation. Compared to WT, *Dcn^−/−^* explants showed accelerated nascent GAG release, both with and without IL-1β (Fig. 3e). For example, by day 6, the cumulative release of nascent GAGs reached 5.1 ± 1.0% and 14.1 ± 1.0% for untreated and IL-1β-treated conditions for WT explants, and 7.6 ± 1.2% and 17.2 ± 1.9% for *Dcn^−/−^* explants, respectively (mean ± 95% CI, *n* ≥ 5). Collectively, these findings demonstrate that loss of decorin accelerates the release of nascent GAGs from the cartilage matrix, highlighting its role in stabilizing newly synthesized aggrecan under both normal and degenerative conditions. In contrast, no genotype or treatment-associated differences were observed in the distribution or release of nascent proteins (Fig. 2b,c and Fig. 3b,f). This indicates that, unlike its role in aggrecan dynamics, decorin does not directly mediate the localization or retention of nascent collagen II molecules.

### 2.3. Exogenous decorin attenuates the loss of nascent GAGs under IL-1β stimulation

Given this stabilizing effect of decorin, we next tested whether supplementing with a supraphysiological level of decorin could attenuate the release of fragmented aggrecan from degenerative cartilage matrix. Immediately after click-labeling, exogenous decorin was added to the culture media at a physiological-like concentration (20 μg/mL)^[24]^ under one of the three regimens: 1) 24 hours prior to an 8-day IL-1β stimulation (pre-treatment), 2) two days after the onset of IL-1β, with media containing decorin refreshed every 2 days (post-treatment), 3) or a combination of both (Fig. 4a). In WT explants, by day 8, decorin pre-treatment significantly reduced IL-1β-induced release of nascent GAGs from 24.5 ± 2.2% to 16.4 ± 1.3% (mean ± 95% CI, *n* = 5, Fig. 4b). In contrast, post-treatment alone (26.9 ± 2.3%) had no appreciable effect, and the combination treatment (17.6 ± 1.1%) did not further enhance the retention compared to pre-treatment alone. These results suggest that increasing decorin levels may slow OA progression by attenuating the loss of fragmented aggrecan. Additionally, the differing outcomes between pre- and post-treatments indicate that decorin is required to integrate into the existing cartilage matrix to exert its stabilizing function. Exposure of free decorin to an IL-1β-induced catabolic environment abolished this effect, possibly due to either degradation of free decorin, the inability of free decorin to affect an already-degraded matrix, or both. Notably, the rescue effects became significant only after a longer culture duration of 8 days, but not within short time frames (≤ 6 days) (Fig. 4b and Fig. S7a). This suggests that decorin-mediated stabilizing effect may be more crucial for preserving cartilage matrix over medium-to-long durations, rather than mitigating acute catabolic responses.

**Figure 4.**
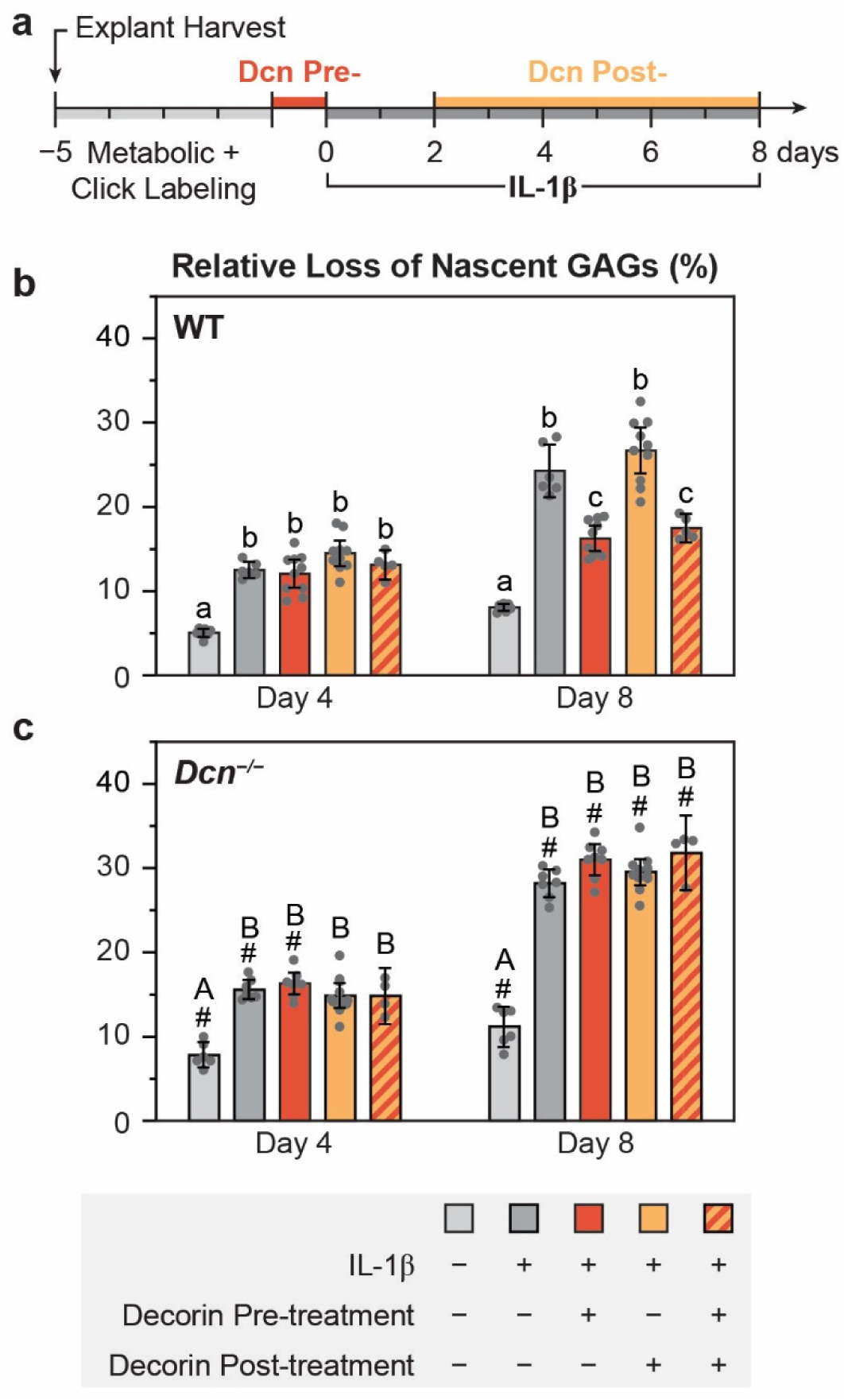
Effects of exogenous decorin on the retention of nascent GAGs in cartilage explants under IL-1β stimulation. a) Schematic of the experimental timeline showing the infiltration of exogenous decorin, click-labeling and IL-1β stimulation in murine femoral head cartilage explants. b,c) Cumulative release of nascent GAGs from b) WT and c) *Dcn^−/−^* cartilage explants at days 4 and 8 under various decorin treatment conditions with IL-1β stimulation. Each data point represents one biological replicate (mean ± 95% CI, *n* ≥ 4). Different letters indicate significant differences among treatments within each genotype and time point. ^#^: *p* < 0.05 between genotypes from the same treatment and time point. A complete list of statistical analysis outcomes for cumulative release of nascent GAGs among different decorin treatment conditions is summarized in Table S9.

In *Dcn^−/−^* explants, neither pre- nor post-treatment of exogenous decorin produced significant effects (Fig. 4c and Fig. S7b). In all treatment conditions, *Dcn^−/−^* cartilage exhibited substantially greater loss of nascent GAGs compared to the WT (Table S9). This unexpected finding may stem from that the infiltrated exogenous decorin did not reach the physiological level of decorin required for its stabilizing function as in WT cartilage. Alternatively, developmental defects in *Dcn^−/−^* cartilage, including reduced aggrecan content and moderate alterations in collagen fibril organization within both the PCM and bulk ECM,^[30–31]^ may impair the integration of exogenous decorin and thus its stabilizing effects. Nevertheless, the pronounced effects observed in WT explants support the potential role of decorin supplementation as a strategy to enhance the preservation of degenerating cartilage matrix.

### 2.4. Decorin stabilizes molecular interactions between aggrecan and collagen II

To elucidate the biophysical mechanisms underlying the impact of decorin on cartilage matrix, we quantified the molecular interaction dynamics among decorin, aggrecan and collagen II using surface plasmon resonance (SPR). We first applied the standard “binary” ligand-analyte protocol to assess real-time binding kinetics between each pair of immobilized ligands and free analytes (Fig. 5a). In this configuration, ligand-analyte association and dissociation kinetics were measured across a series of decreasing analyte concentrations. Monte Carlo simulation^[48]^ was applied to analyze the resulting sensorgrams, identify the appropriate interaction model (Table 2), and then calculate the predicted equilibrium dissociation constant, *K_D_*, as a measure of binding affinity (Fig. 5b).

**Figure 5.**
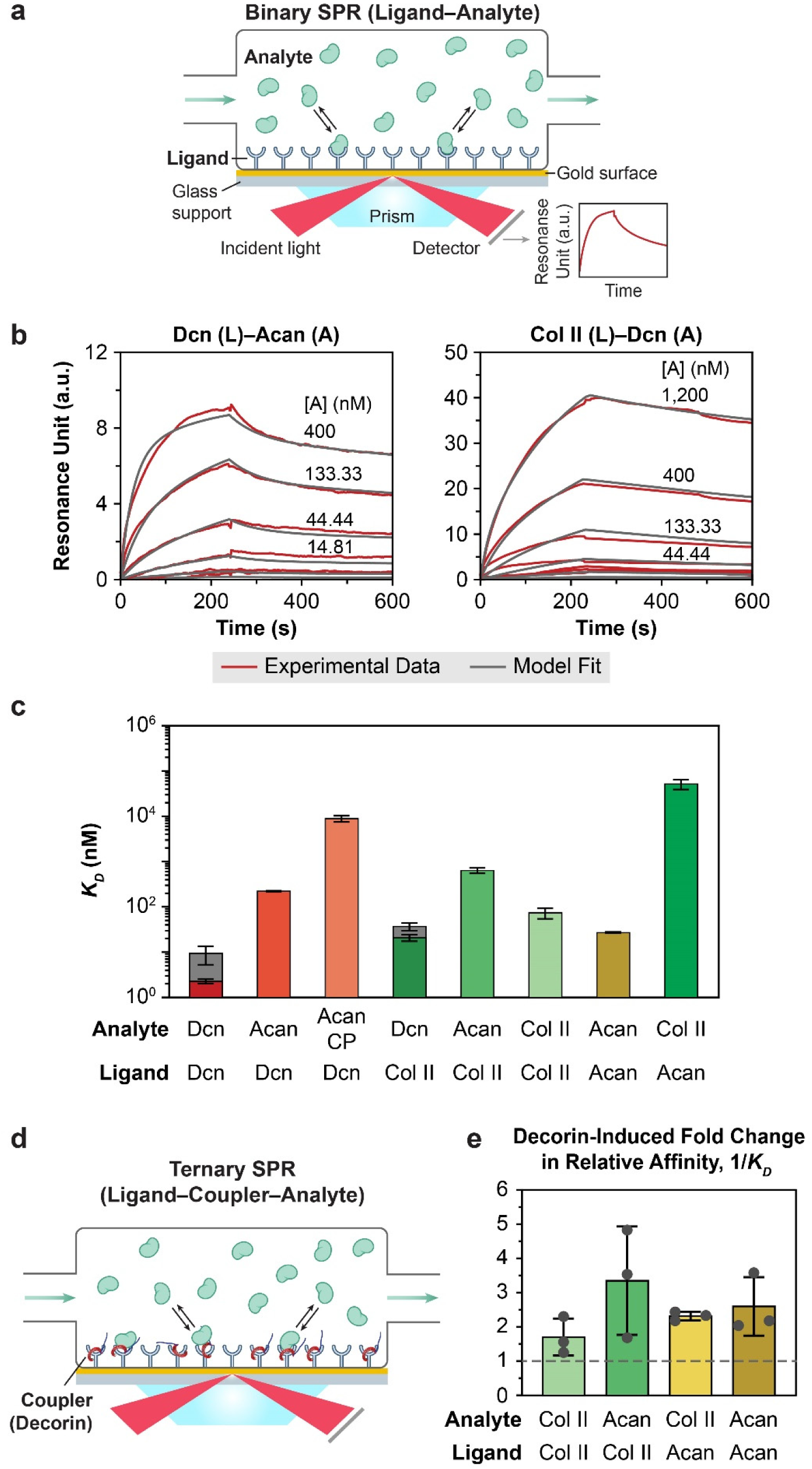
Surface plasmon resonance (SPR) quantification of binding affinities between decorin and major cartilage matrix constituents, highlighting the role of decorin in integrating collagen II and aggrecan. a) Schematic illustration of the binary SPR setup for assessing ligand-analyte (*L*-*A*) binding kinetics. b) Representative sensorgrams for decorin (*L*)-aggrecan (*A*) and collagen II-decorin interactions showing raw experimental data at decreasing analyte concentrations (red) and best fit models (black), derived via Monte Carlo simulation using the two-step maturation and heterogeneous ligand binding models, respectively. c) Semi-log plots of the equilibrium dissociation constants (*K_D_*) comparing binding affinities across all bi-element pairs (mean ± SD derived from 50 cycles of Monte Carlo model fits to four independent replicates per group, Table 2). Values represent *K_D_*_1_ for all binding pairs, except for decorin-decorin and collagen II-decorin binding, for which, the range of independent *K_D_*_1_ and *K_D_*_2_ are shown as dark grey zones from the heterogeneous ligand model (Table 3). d) Schematic illustration of the “ternary” SPR protocol for quantifying the effects of decorin as a molecular coupler on ligand-analyte binding. e) Decorin-induced fold changes in the relative binding affinity, 1/*K_D_*, for all tested conditions. Each data point represents one independent replicate (mean ± SD, *n* = 3). A complete list of detailed experimental outcomes for each replicate is summarized in Table S10.

**Table 2.**
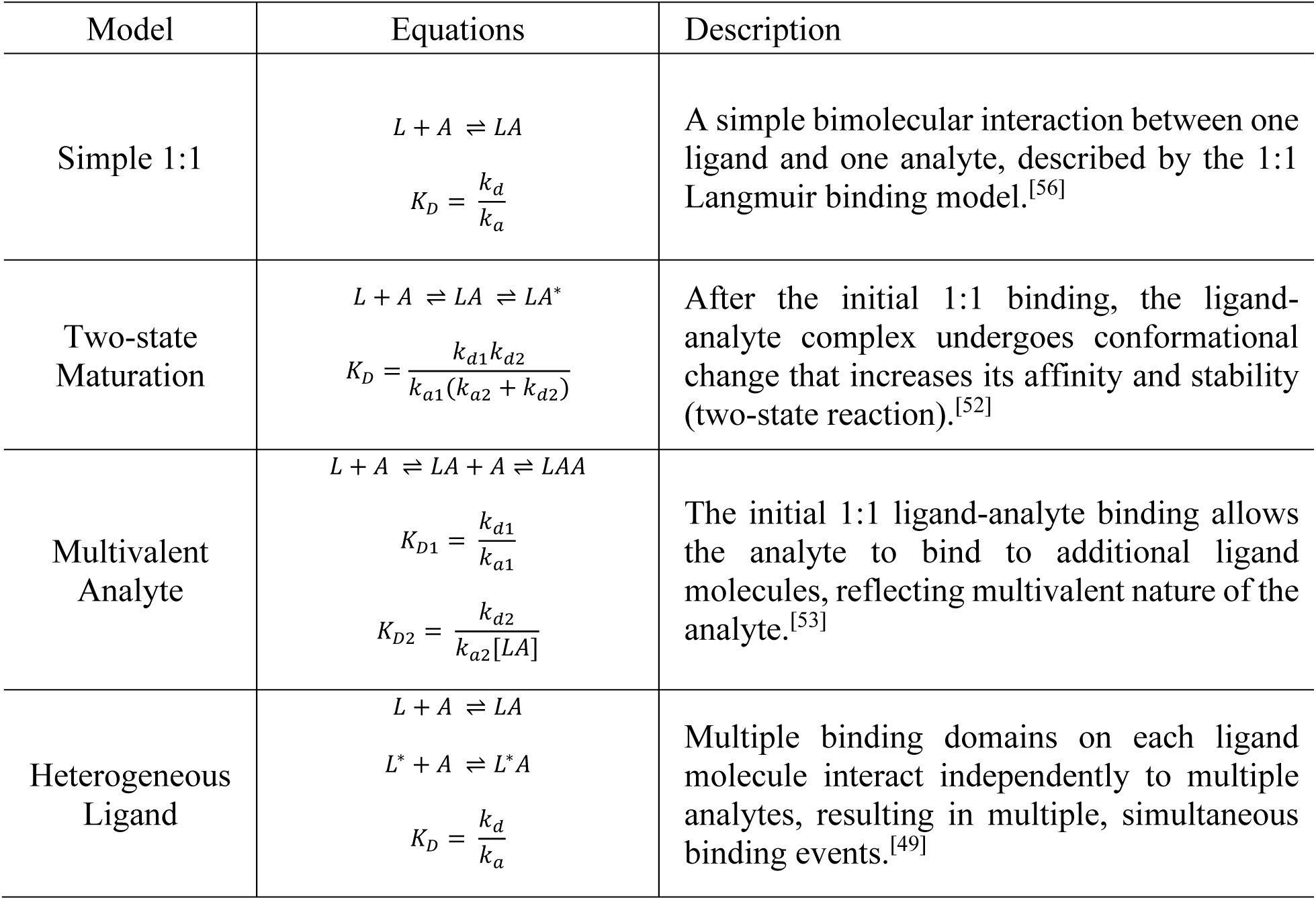
Glossary of molecular interaction kinetics models describing ligand (*L*)-analyte (*A*) binding measured by surface plasmon resonance (SPR).

**Table 3.**
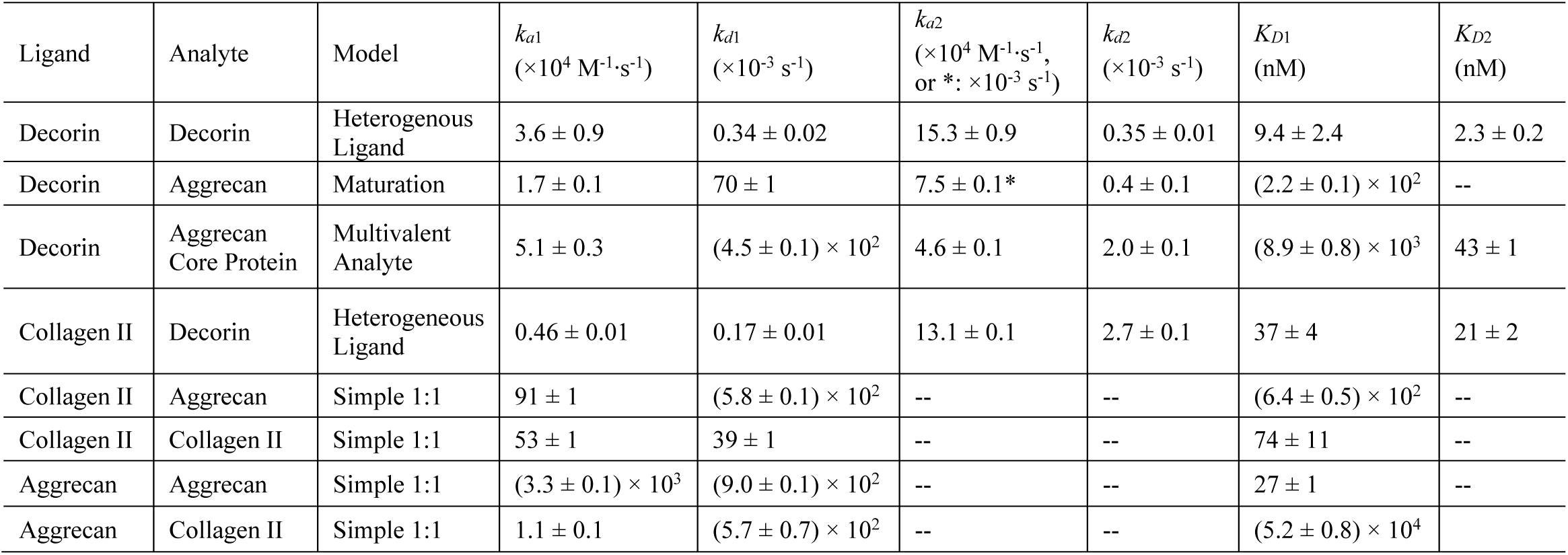
List of predicted models describing molecular interaction kinetics models and parameters for binary surface plasmon resonance (SPR) setup, including the association and dissociation rate constants (*k_a_* and *k_d_*), equilibrium dissociation constant (*K_D_*), shown as mean ± SD from 50 Monte Carlo simulation cycles fitted with sensorgrams from *n* = 3 replicates for each group.

| Ligand | Analyte | Model | $k_{a1}$<br>($\times 10^4 \text{ M}^{-1} \cdot \text{s}^{-1}$) | $k_{d1}$<br>($\times 10^{-3} \text{ s}^{-1}$) | $k_{a2}$<br>($\times 10^4 \text{ M}^{-1} \cdot \text{s}^{-1}$,<br>or *: $\times 10^{-3} \text{ s}^{-1}$) | $k_{d2}$<br>($\times 10^{-3} \text{ s}^{-1}$) | $K_{D1}$<br>(nM) | $K_{D2}$<br>(nM) |
| --- | --- | --- | --- | --- | --- | --- | --- | --- |
| Decorin | Decorin | Heterogenous Ligand | $3.6 \pm 0.9$ | $0.34 \pm 0.02$ | $15.3 \pm 0.9$ | $0.35 \pm 0.01$ | $9.4 \pm 2.4$ | $2.3 \pm 0.2$ |
| Decorin | Aggrecan | Maturation | $1.7 \pm 0.1$ | $70 \pm 1$ | $7.5 \pm 0.1^*$ | $0.4 \pm 0.1$ | $(2.2 \pm 0.1) \times 10^2$ | -- |
| Decorin | Aggrecan Core Protein | Multivalent Analyte | $5.1 \pm 0.3$ | $(4.5 \pm 0.1) \times 10^2$ | $4.6 \pm 0.1$ | $2.0 \pm 0.1$ | $(8.9 \pm 0.8) \times 10^3$ | $43 \pm 1$ |
| Collagen II | Decorin | Heterogeneous Ligand | $0.46 \pm 0.01$ | $0.17 \pm 0.01$ | $13.1 \pm 0.1$ | $2.7 \pm 0.1$ | $37 \pm 4$ | $21 \pm 2$ |
| Collagen II | Aggrecan | Simple 1:1 | $91 \pm 1$ | $(5.8 \pm 0.1) \times 10^2$ | -- | -- | $(6.4 \pm 0.5) \times 10^2$ | -- |
| Collagen II | Collagen II | Simple 1:1 | $53 \pm 1$ | $39 \pm 1$ | -- | -- | $74 \pm 11$ | -- |
| Aggrecan | Aggrecan | Simple 1:1 | $(3.3 \pm 0.1) \times 10^3$ | $(9.0 \pm 0.1) \times 10^2$ | -- | -- | $27 \pm 1$ | -- |
| Aggrecan | Collagen II | Simple 1:1 | $1.1 \pm 0.1$ | $(5.7 \pm 0.7) \times 10^2$ | -- | -- | $(5.2 \pm 0.8) \times 10^4$ | |

We found that collagen II-decorin and decorin-decorin bindings were best described by the heterogeneous ligand model,^[49]^ consistent with previous studies showing multiple decorin-binding sites on collagen II^[50]^ and multiple leucine-rich repeats (LRRs) dimerization domains along the concave surface of decorin core protein.^[51]^ Based on this model, we derived two parallel binding kinetics parameters with different *K_D_*’s for each pair. Decorin-decorin interaction showed the highest binding affinity, as reflected by the lowest *K_D_* values compared to other tested pairs (Fig. 5c and Table 3), supporting the formation of stable decorin-decorin dimer complexes. In comparison, collagen II-decorin interaction exhibited either slower association rate (*k_a_*_1_) or faster dissociation rate (*k_d_*_2_), yielding higher magnitudes of both *K_D_*_1_ and *K_D_*_2_ indicative of moderately lower binding stability.

Decorin-aggrecan interaction was described by the two-step maturation model,^[52]^ suggesting that the ligand-analyte complex formation was followed by conformational changes to further stabilize the interaction. This model yielded association (*k_a_*_1_) and maturation rates (*k_a_*_2_) of the same order as decorin-decorin *k_a_*_1_, but a much faster dissociation rate *k_d_*_1_, resulting in considerably lower binding affinity compared to decorin-decorin and collagen II-decorin. However, decorin-aggrecan core protein interaction was best described by the multivalent analyte model, consistent with the initial binding inducing conformational changes of aggrecan core protein that promote additional tandem interactions with decorin.^[53]^ This interaction could arise from the globular domains along aggrecan core protein that could potentially provide multiple conformation-dependent binding sites.^[54–55]^ Based on this model, we noted approximately 3× faster association rate (*k_a_*_1_) but ≈ 60× faster dissociation rate (*k_d_*_1_), resulting in a ≈ 40× higher *K_D_*_1_ indicative of much lower binding affinity compared to decorin-intact aggrecan interaction. These results suggested that while aggrecan core protein initiates decorin-aggrecan binding, GAG-mediated conformational changes are required to stabilize their interactions. Taken together, our results support that decorin directly interacts with decorin, collagen II and aggrecan, with the highest affinity for decorin-decorin self-association, followed by collagen II-decorin interaction.

To compare decorin molecular interactions with those between aggrecan and collagen II, we applied the simple 1:1 Langmuir model^[56]^ to collagen II-collagen II, aggrecan-aggrecan and collagen II-aggrecan sensorgrams to focus on their binding affinity rather than detailed molecular kinetics. We found that collagen II-collagen II and aggrecan-aggrecan interactions both showed similar orders of *K_D_*s compared to that of collagen II-decorin. Collagen II-aggrecan and aggrecan-collagen II interactions, on the other hand, have higher *K_D_*s indicative of lower binding affinity, suggesting that direct association between these two major constituents likely does not have a major contribution to cartilage matrix assembly. Here, the much higher *K_D_* observed for aggrecan (ligand)-collagen II (analyte) compared to collagen II (ligand)-aggrecan (analyte) was likely associated with the high long-range electrical double layer (EDL) repulsion from the negatively charged aggrecan substrate that inhibits protein binding. The same effect likely contributed to the consistently faster *k_d_*_1_’s observed for all interactions involving aggrecan, except for that of decorin-aggrecan binding.

Lastly, given the active interactions among decorin, collagen II and aggrecan, we queried how decorin impacts the interaction affinities between collagen II and aggrecan. We applied a “ternary” complex SPR setup,^[57]^ in which decorin was supplied as a “coupler” to bind to the ligand and potentially mediate ligand-analyte interactions between collagen II and aggrecan (Fig. 5d). We then applied the same simple 1:1 model via Scrubber to individual sets of sensorgrams for collagen II-collagen II, aggrecan-aggrecan and collagen II-aggrecan to investigate changes in relative binding affinity with versus without decorin as a coupler, rather than elucidating the molecular kinetics by Monte Carlo simulation. Despite the expected substantial variations in the extracted apparent *K_D_*’s due to sample-to-sample variations in ligand heterogeneity and ligand mass transport^[49]^ (Table S10), we noted that addition of decorin as a coupler consistently enhanced all measured aggrecan-collagen II interactions (Fig. 5e and Table S10), with fold changes in 1/*K_D_* ranging from 1.7 ± 0.5 to 3.4 ± 1.4 (*n* = 3, mean ± SD, Fig. 5e). Therefore, these findings support that by directly binding to collagen II and aggrecan, decorin-mediated interactions strengthen the integration of aggrecan and collagen II through stabilizing the formation of their molecular complexes.

### 2.5. Decorin strengthens the collagen II fibril network

Given the higher binding affinity of decorin-collagen II (Fig. 5c) and the stabilizing effect of decorin on collagen II-collagen II interactions (Fig. 5e), we next investigated whether decorin-collagen II association also modulates the mechanical properties of the collagen II fibril network, which are essential for confining resident aggrecan.^[8]^ We enzymatically removed all proteoglycans and other non-fibrillar constituents from femoral condyle cartilage, and infiltrated the GAG-depleted cartilage with 20 µg/mL exogenous bovine decorin. GAG removal and decorin infiltration were validated via histology and immunofluorescence (IF) imaging (Fig. 6a,b). Applying AFM-nanoindentation, we quantified the effects of decorin infiltration on the indentation modulus of untreated and GAG-depleted cartilage for both genotypes (Fig. 6c). In WT cartilage, GAG depletion significantly reduced the modulus by 71 ± 6% (0.40 ± 0.08 MPa for untreated and 0.12 ± 0.02 MPa for GAG-depleted, *p* < 0.001, *n* ≥ 12), reaffirming the expected contribution of aggrecan and its negative charges to cartilage mechanical properties.^[58]^ Notably, infiltration of decorin significantly increased the modulus of GAG-depleted cartilage to 0.25 ± 0.07 MPa (*p* < 0.05, *n* = 8). As the decorin infiltration had minimal effect on restoring CS-GAG content and its fixed charges (Fig. 6a), these results highlight the role of decorin in strengthening the collagen II network. This augmenting effect, however, was not observed in untreated tissue, likely because the aggrecan-endowed osmotic swelling masked the direct contribution from decorin. Furthermore, exogenous decorin did not increase the modulus of *Dcn^−/−^* cartilage, either in untreated or GAG-depleted conditions (Fig. 6c). This suggests that exogenous decorin cannot reinforce the altered collagen II fibril network formed in the absence of endogenous decorin, consistent with the lack of rescue effect of IL-1β-induced nascent GAG loss from *Dcn^−/−^* explants (Fig. 4c).

**Figure 6.**
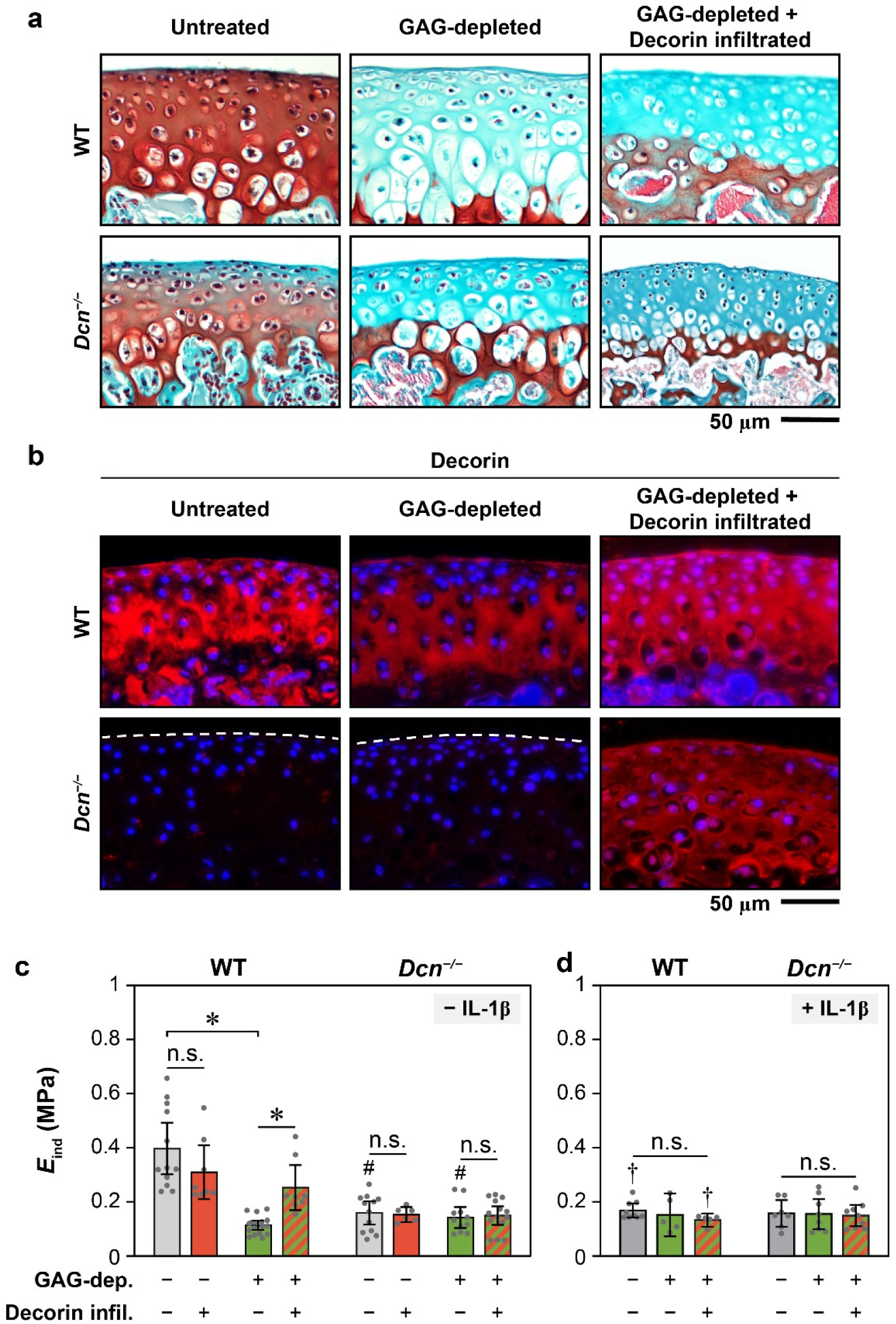
Effects of exogenous decorin infiltration on the modulus of murine cartilage and its collagen II fibrillar network. a,b) Representative images of a) Safranin-O/Fast Green histology and b) decorin immunofluorescence (IF) staining of 3-week-old wild-type (WT) and decorin-null (*Dcn^−/−^*) femoral condylar cartilage under untreated, GAG-depleted and GAG-depleted with decorin infiltration conditions. Images confirm effective GAG removal and illustrate decorin infiltration throughout cartilage without appreciable changes in GAG staining. c,d) Indentation modulus, *E*_ind_, of medial femoral condylar cartilage from 3-week-old WT and *Dcn^−/−^* mice measured via AFM-nanoindentation following the GAG depletion and decorin infiltration in c) freshly harvested tissues without IL-1β stimulation and d) IL-1β-stimulated tissues. Each data point represents the average value of one animal (mean ± 95% CI, *n* ≥ 6). *: *p* < 0.05 between treatment groups within the same genotype, ^#^: *p* > 0.05 between genotypes for the same treatment, ^†^: *p* < 0.05 between with versus without IL-1β for the same genotype, n.s.: not significant. A complete list of statistical analysis outcomes for *E*_ind_ among different treatment conditions is summarized in Table S11.

To validate that the strengthening effect of decorin depends on an intact collagen II network, we repeated the AFM-nanoindentation on IL-1β-treated cartilage followed by GAG-depletion and decorin infiltration. For both genotypes, decorin infiltration did not have a significant impact on the modulus of these IL-1β-treated, GAG-depleted tissues (Fig. 6d). In addition, for WT cartilage treated with IL-1β, as well as *Dcn^−/−^* cartilage with or without IL-1β, GAG depletion did not significantly reduce the modulus, indicating that in these degenerative tissues, direct contribution of aggrecan and its GAGs to tissue modulus is diminished due to matrix damage. Consequently, the ability of decorin to strengthen the matrix is lost when the collagen fibril network integrity is compromised.

## 3. Discussion

This study identifies decorin as an indispensable constituent in regulating the dynamic turnover of cartilage matrix, particularly in retaining newly synthesized aggrecan. In cartilage, the primary mechanism of aggrecan assembly is its formation of aggregates with HA^[6]^ through link protein-assisted binding^[7]^ at its G1-globular domain. However, this mechanism alone does not fully account for the retention of a substantial portion of G1-deficient aggrecan fragments^[59]^ that result from normal homeostasis.^[60–61]^ Our results establish a crucial role for decorin in stabilizing nascent aggrecan within native cartilage matrix. This role complements the aggrecan-HA assembly mechanism, contributing to the retention of aggrecan under both normal homeostatic and disease-associated catabolic conditions (Fig. 3e). On the other hand, despite its established role in binding to collagen triple helix,^[62]^ we show that decorin does not directly regulate the distribution or retention of nascent collagen in cartilage. While decorin may influence the structure or mechanics of collagen II fibril network, the retention of collagen II triple helices and their assembly into fibrils are likely governed by other regulatory collagens, such as types III, IX and XI.^[63–64]^

We attribute this stabilizing function of decorin to two biophysical mechanisms (Fig. 7). First, through directly interacting with aggrecan and collagen II, decorin provides molecular linkages to form supramolecular networks that enhance the integration of aggrecan aggregates and collagen II fibrillar network (Fig. 7). Our previous molecular force spectroscopy studies showed that aggrecan exhibits molecular adhesions with other aggrecan molecules and collagen II fibrils, contributing to its matrix retention.^[65–66]^ Although decorin-aggrecan binding demonstrates a lower affinity compared to collagen II-decorin and decorin-decorin (higher *K_D_*, Fig. 5c), decorin may still endow additional molecular interactions to reinforce the formation of collagen II-aggrecan complex. This is supported by our ternary SPR outcomes confirming the ability of decorin to strengthen aggrecan-aggrecan and aggrecan-collagen II interactions (Fig. 5e), as well as our earlier findings showing decorin increases the aggrecan-aggrecan and aggrecan-collagen II molecular adhesion.^[30]^ Second, by connecting collagen II molecules (Fig. 5c,f), decorin strengthens the collagen II fibrillar network (Fig. 7). Given the comparable *K_D_*’s between decorin-decorin and collagen II-decorin, decorin can potentially fortify the collagen II network through connecting adjacent fibrils, likely through its monomer form,^[67]^ as supported by the strengthening effect on collagen II-collagen II interactions (Fig. 5e). At the fibril level, this effect was further illustrated by the increased modulus of decorin-infiltrated collagen II fibril network of WT cartilage (Fig. 6c). Given the essential role of collagen II fibril network in confining densely packed aggrecan,^[8]^ this strengthening effect of decorin enhances resistance to the diffusive loss of aggrecan driven by osmotic pressure, either in its aggregated or monomer forms. Collectively, these decorin-endowed interactions work in synergy to increase aggrecan-collagen II integration, contributing to the overall ECM integrity.

**Figure 7.**
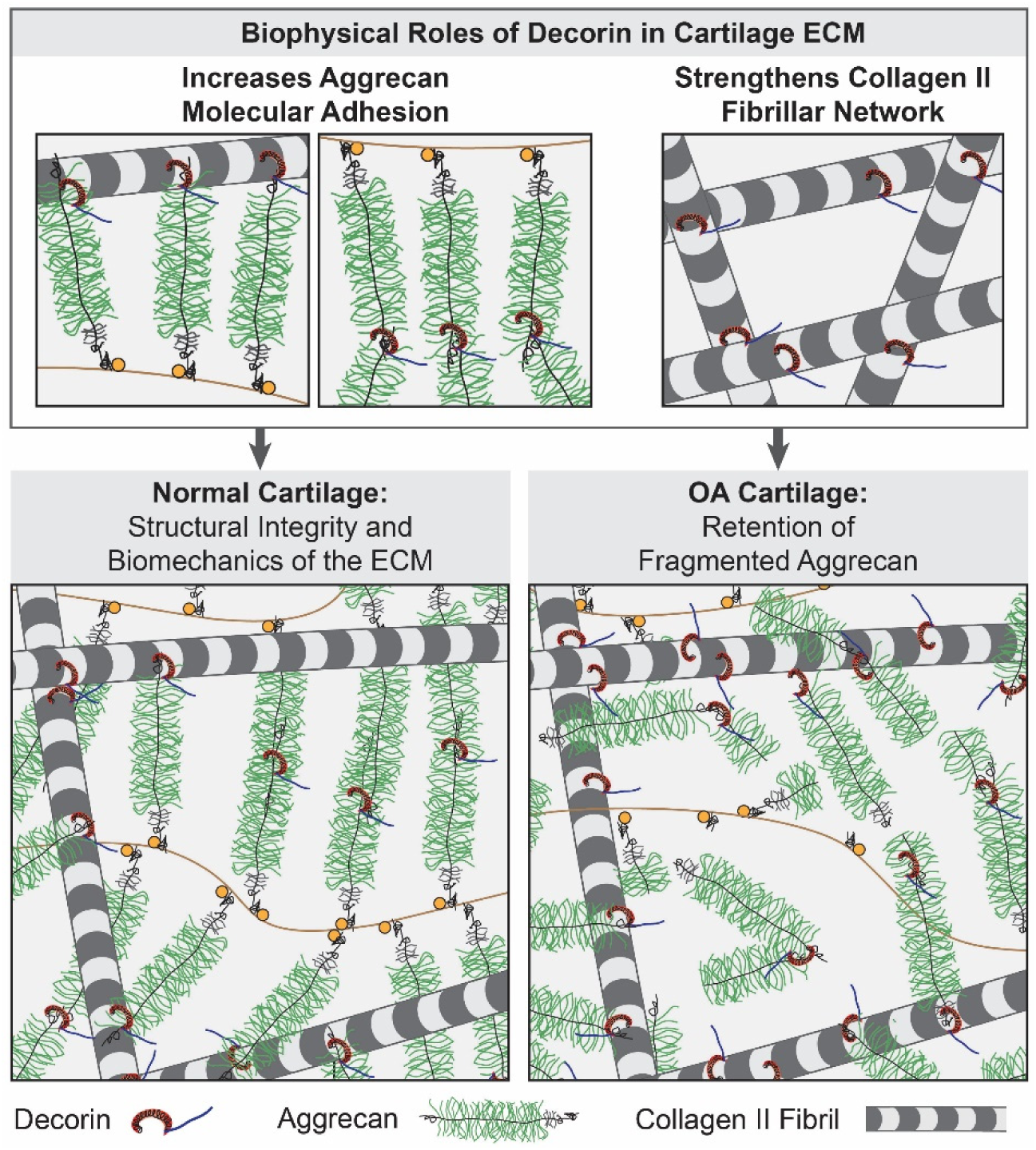
Schematic illustration highlighting the biophysical roles of decorin in maintaining cartilage matrix integrity. Decorin interacts with both collagen II fibrils and aggrecan to increase the molecular adhesion essential for collagen II-aggrecan integration. In addition, by binding to collagen II fibrils, decorin strengthens the fibrillar network, reinforcing its role in entrapping the highly compressed aggrecan within the porous network. This decorin-mediated mechanism is vital for aggrecan retention during normal matrix turnover and delays the loss of fragmented aggrecan under degenerative conditions. In the schematics, the packing densities of collagen II fibrils and aggrecan aggregates are reduced to increase clarity.

Notably, this decorin-mediated collagen II-aggrecan integration does not affect the preferential localization of nascent aggrecan in the PCM. We found that the peak concentration of nascent aggrecan is ≈ 900 nm from the cell surface (*d*_peak_, Fig. 3d), which is ≈ 2× the contour length of full-length aggrecan core protein (≈ 350-550 nm).^[59,68]^ This localization is likely driven by specific interactions at the chondrocyte surface, such as coordination between aggrecan-HA^[6]^ and HA-CD44^[69]^ associations, or other interactions involving PCM-exclusive molecules.^[70]^ The distinct structural features of PCM, including thinner collagen fibrils and smaller inter-fibrillar spaces,^[31]^ may also contribute to the transient retention of nascent aggrecan. Since decorin is present in both the PCM and bulk ECM (e.g., Fig. 6b), our findings suggest that its stabilizing role is independent of aggrecan-HA association and does not regulate aggrecan localization. Therefore, decorin functions as a stabilizer, rather than a primary driver of aggrecan assembly in cartilage matrix.

We further clarify that the primary role of decorin in cartilage is to regulate the matrix integrity, rather than directly altering chondrocyte signaling. Despite pronounced matrix defects in postnatal *Dcn^−/−^* cartilage,^[30]^ RNA-sequencing results revealed no significant transcriptomic changes, including in TGF-β signaling (Fig. 1b-d) and matrisome gene expression (Fig. S6). This suggests that decorin does not directly influence chondrocyte fate or mechanosensitive signaling *in vivo*. The previously reported biological functions of decorin^[36]^ are likely dependent on tissue types and disease states, and are not prominently active in cartilage. This absence of gene expression changes also contrasts with altered intracellular calcium signaling observed in 3-month-old *Dcn^−/−^*cartilage,^[31]^ suggesting that chondrocytes retain a certain degree of innate resilience to microenvironment perturbations. Similar resilience to matrix perturbation was observed in the meniscus of *Col5a1^+/−^* mice, a model for classical Ehlers-Danlos Syndrome (cEDS),^[71]^ illustrating an inherently robust mechanosensing framework to preserve normal homeostasis in various biological systems *in vivo*.^[72]^ However, the disrupted matrix integrity and cell mechanosensing may indicate higher susceptibility to disease initiation, as supported by the more pronounced OA severity observed in *Dcn^−/−^* and induced decorin knockout mice following the DMM surgery.^[33–34]^

Our results support decorin modulation as a promising OA intervention strategy. In early OA, aggrecan fragmentation and loss^[17–18]^ disrupt chondrocyte mechanotransduction and leads to irreversible cartilage damage. Decorin-strengthened matrix delays the loss of fragmented aggrecan (Fig. 4), supporting the potential of elevating decorin to supraphysiological levels to enhance cartilage preservation. However, multiple factors warrant consideration. The absence of post-IL-1β rescue may be due to decorin degradation or failed decorin-matrix integration in the catabolic environment. Also, the decorin-stiffening effect requires an intact collagen II fibril network and is not observed in IL-1β-treated WT cartilage or *Dcn^−/−^* tissue (Fig. 6c,d). This aligns with the lack of decorin rescue effects under these conditions (Fig. 4c). Nevertheless, these findings do not preclude the therapeutic potential of decorin. In OA, cartilage degradation usually initiates at the sites of injury or those subjected to excessive weight bearing, and the degree of matrix damage is heterogeneous.^[73]^ Targeted decorin delivery to moderately damaged regions may preserve cartilage and prevent further degeneration. Gene therapy to enhance endogenous decorin expression may offer additional advantages by promoting direct integration of endogenous decorin with the PCM. Unlike biglycan,^[29]^ decorin fragmentation does not appear to exacerbate chondrocyte catabolic signaling, supporting the safety of this strategy. Our ongoing work explores the use of decorin-based biomaterials such as synthetic decorin-mimetics,^[74]^ recombinant decorin,^[75]^ or decorin-targeting gene therapies^[76]^ to promote cartilage matrix preservation.

Finally, this newly discovered role of decorin in aggrecan-collagen II integration provides new insights relevant to other diseases involving aberrant aggrecan turnover and degradation, including intervertebral disc (IVD) degeneration,^[77]^ temporomandibular joint (TMJ) disorder,^[78]^ rheumatoid and psoriatic arthritis,^[79]^ as well as *ACAN* mutation-associated genetic disorders.^[80]^ Given its canonical function in regulating collagen I fibrillogenesis,^[81]^ decorin also acts as an anti-fibrotic agent in conditions marked by excessive deposition and aberrant assembly of collagen I.^[82]^ Based on our findings that decorin strengthens collagen II fibril network (Fig. 7), this work supports further investigation of decorin activities in mitigating diseases involving abnormal collagen I remodeling, such as wound healing^[83]^ and solid tumors.^[84]^ On the technical front, this study showcases the utility of bio-orthogonal click-labeling as a powerful tool to uncover the turnover and dynamics of nascent matrix proteoglycans and proteins, and to distinguish the activities of newly synthesized versus existing matrix molecules. When combined with imaging analysis and other molecular modalities such as SPR and AFM-nanomechanical tests, this approach enables molecular dissection of regulatory matrix molecules such as decorin and offers a versatile platform to study ECM turnover and remodeling.

## 4. Conclusion

This study highlights a crucial molecular interaction axis of collagen II-decorin-aggrecan in regulating cartilage matrix integrity. Despite being a quantitatively minor constituent, decorin plays a pivotal role in promoting collagen II-aggrecan integration by stabilizing their molecular interactions and strengthening the collagen II fibril network that entraps aggrecan. These results provide new molecular-level insights into the dynamic assembly and turnover of cartilage matrix, extending beyond the established paradigm of aggrecan-HA assembly. Importantly, elevating decorin to a supraphysiological level effectively attenuates the loss of nascent aggrecan caused under IL-1β-induced catabolic conditions, underscoring its potential as a molecular therapeutic for preserving degenerating cartilage. These results also shed light on the broader significance of decorin in the assembly, degeneration and repair of other cartilaginous and fibrocartilaginous tissues enriched in aggrecan, such as intervertebral disc and trachea.

## 5. Experimental Section

### Animal model

Decorin-null (*Dcn^−/−^*) mice in the C57BL/6 strain were generated as previously described^[41]^ and housed in the Calhoun animal facility at Drexel University. Tissues were harvested from 3-week-old WT and *Dcn^−/−^* mice from littermates of *Dcn^+/−^* breeders. Both male and female mice were included, as we did not observe sex-associated variations in *Dcn^−/−^* cartilage phenotype.^[30]^ All animal work was approved by the Institutional Animal Care and Use Committee (IACUC) at Drexel University (Protocol #LA-22-058).

### Bulk RNA-sequencing

Total RNA was extracted from freshly dissected femoral heads of 1-month-old WT and *Dcn^−/−^* mice using TRI-reagent (T9424, Sigma).^[30]^ A total of four femoral heads from two siblings were pooled as one biological replicate for each genotype (*n* = 3 replicates/genotype) to obtain sufficient RNA. RNA sequencing was performed by Eukaryotic Transcriptome (BGI America). In brief, raw sequencing data underwent quality control and filtering to remove low-quality reads prior to genomic alignment with HISAT.^[85]^ Following alignment, transcript prediction was performed with StringTie^[86]^ and Cuffcompare,^[87]^ with coding potential assessed with Coding Potential Calculator (CPC).^[88]^ To evaluate sequence variation across samples, the Genome Analysis Toolkit (GATK)^[89]^ was applied to detect single-nucleotide polymorphisms (SNPs) and insertions/deletions (INDELs). Following this, clean reads were mapped to reference transcripts using Bowtie2.^[90]^ Finally, DESeq2^[91]^ was used to identify differentially expressed genes (DEGs) as those with adjusted *p*-value < 0.05 from the Wald test followed by Benjamini-Hochberg correction for family-wise type I errors. To perform principal component analysis (PCA), normalized read counts for each gene across biological replicates were used as inputs, with gene names serving as identifiers, using the PCA function in the online tool SRplot. Normalized gene counts from available KEGG canonical^[37,42]^ and non-canonical^[43]^ TGF-β signaling pathways (KEGG: mmu04350), Gene Ontology (GO) Consortium datasets related to decorin binding, including EGFR (GO: 0007173),^[44]^ VEGF (GO: 0038084),^[45]^ and Met-mediated HGFR (GO: 0048012),^[46]^ as well as top 40 relatively abundant genes from the Matrisome Database^[92]^ were used to generate bidirectionally clustered heatmaps using the online tool SRplot (Bioinformatics).^[93]^

### Cartilage explant culture and bio-orthogonal click-labeling

Femoral head cartilage explants were isolated from 3-week-old mice and pre-cultured for 2 days in chondrogenic DMEM for equilibration, following our established procedure.^[33]^ To label newly synthesized GAGs and proteins, explants were cultured in chondrogenic DMEM supplemented with either 30 μM N-azidoacetylgalactosamine-tetraacylated (GAL, CCT-1086, VectorLabs) or 30 μM L-Azidohomoalanine (AHA, CCT-1066, VectorLabs) for 3 days, respectively.^[40]^ Explants were then cultured in DMEM with 30 μM of AZDye 488 DBCO (MB488, CCT-1278, VectorLabs) for 2 hours to fluorescently tag newly synthesized GAGs or proteins via click chemistry and subsequently washed overnight to remove unreacted dye. Finally, they were cultured in chondrogenic DMEM with or without 10 ng/mL interleukin-1β (IL-1β, 211-11B, Peprotech)^[94]^ for 6 days to induce chondrocyte catabolism, with media refreshed and collected every 2 days (Fig. 2a). On day 6, explants were digested with 2% papain (76216, MilliporeSigma). Fluorescent intensity from both the media and digested explants was measured at excitation at 488 nm and emission at 530 nm to quantify the amounts of nascent GAGs/proteins released to the media versus those retained in the explants.

To evaluate the rescue effects of decorin, exogenous decorin (D8428, MilliporeSigma) was introduced under three scenarios (Fig. 4a). First, for decorin pre-treatment, immediately after click-labeling, explants were cultured with 20 µg/mL decorin for 24 hours before IL-1β stimulation for 8 days, with media refreshed and collected every 2 days. Second, for decorin post-treatment, decorin was added on day 2 of IL-1β stimulation and was refreshed every two days up to day 8. Third, explants were subjected to both pre- and post-treatment of decorin. In each case, the nascent GAGs/proteins released to the media or retained in the explants were quantified.

### Confocal imaging and data analysis

To assess spatial distributions of nascent GAGs/proteins, explants were imaged using a Zeiss LSM700 confocal microscope (63×, emissions at 358, 488 and 657 nm) immediately before IL-1β onset and on days 2, 4 and 6 of IL-1β culture. Cell viability was monitored by staining with 5 μg/mL Hoechst 33342 (H3570, Invitrogen) for cell nuclei and 4 µM ethidium homodimer I (E-1903, MilliporeSigma) for dead cells prior to imaging.

A custom MATLAB image analysis program was developed to quantify the intensity and spatial distribution of fluorescent signals in the explants. High-resolution images were read for intensity per pixel, and co-planar chondrocytes were contoured. ROIs (≈ 100 × 100 µm^2^) containing the intracellular spaces for at least 6 chondrocytes were selected. Within each ROI, the maps of both fluorescent signal intensity and the Euclidean distance to the nearest chondrocyte surface, *d*_Matrix-to-Cell_, per pixel (Fig. 2d) were assessed. We calculated the average intensity, *I*_mean_, for each ROI. Furthermore, we adjusted the intensity signal values to correspond to regular 10 nm intervals of *d*_Matrix-to-Cell_ (Fig. 2d) and calculated the average and interquartile range (IQR) of the intensity as a function of *d*_Matrix-to-Cell_ (solid black line and dark red region in Fig. 2d, respectively). We then computed the spatial parameters including the peak intensity, *I*_peak_, the distance corresponding to peak intensity, *d*_peak_, and the half-decay distance corresponding to the mid-point between peak and baseline intensities, *d*_1/2_ (Fig. 2d,e and Table 1). For AHA images, given the absence of clear signal peaks (Fig. 2f), we only calculated *I*_mean_ for each ROI.

### Histology and Immunofluorescence (IF) Imaging

Femoral condyles from 3-week-old WT and *Dcn^−/−^* mice were enzymatically treated with 0.5 U/mL chondroitinase ABC and 500 U/mL hyaluronidase to remove GAGs, followed by 24 hours incubation in 1× PBS with 20 µg/mL exogenous bovine decorin. Histology and immunofluorescence (IF) imaging were performed on untreated, GAG-depleted, as well as GAG-depleted samples following decorin infiltration. Samples were fixed in 4% paraformaldehyde, decalcified in 10% EDTA for 3 weeks, and embedded in paraffin. Serial 6 μm-thick sagittal sections were cut across the condyle. Sections were stained with Safranin-O/Fast Green and imaged in brightfield using a Leica DM4000 B microscope (Leica) to assess sGAG distribution and confirm its removal. For IF imaging, additional sections were incubated with the primary antibody for decorin (LF-114 [ENH019-FP], Kerafast, 1:100 dilution) in 1% bovine serum albumin (BSA) overnight at 4 °C, and incubated with the secondary antibody (A11008, Invitrogen) for 1 hour at room temperature, before mounted with DAPI Fluoromount-G (0100-20, SouthernBiotech) as per the established procedure.^[31]^ Fluorescence images were taken using a Leica DMI-6000B microscope with 358 nm emission filter and a 488 nm fluorescent lamp at 50% power and 300 ms exposure time in a 16-bit, 1,392 × 1,040 imaging array format.

### Surface plasmon resonance (SPR) and data analysis

Two-component, or “binary” SPR setup (BIAcore S200) was applied to quantify the molecular interaction and binding kinetics among decorin, aggrecan and collagen II, following the established procedure.^[95–97]^ Aggrecan core protein was prepared by enzymatic treatment of intact aggrecan (A1960, MilliporeSigma) with a cocktail of 0.5 U/mL chondroitinase ABC (C3667, MilliporeSigma) and 500 U/mL hyaluronidase (37326-33-3, MP Biomedicals) in 1× PBS for 24 hrs. Ligand molecules, decorin and collagen II (C1188, MilliporeSigma), were filtered and immobilized onto a carboxymethylated polyethylene glycol (CM-PEG)-based flat sensor at pH 7.4 (PEG chip 29239810, Cytiva and XanTec), at a concentration of 50-200 μg/mL in 1× PBS (pH 7.4) via coupling with freshly mixed 1:1 ratio of 200 mM 1-ethyl-3-(3-(dimethylamino) propyl) carbodiimide (EDC) and 50 mM N-hydroxysuccinamide (NHS). Immobilization was established at ligand response units (RU) between 90-200, and PEG-based sensors were selected for their higher efficiency at minimizing non-specific adsorption than standard gold surfaces.^[98]^ Due to the net negatively charged nature of decorin and aggrecan, PBS-P (PBS with 0.05% polysorbate, P6585, MilliporeSigma), rather than the standard 1 M ethanolamine, was injected for 10-15 minutes to hydrolyze unreacted EDC-NHS sites.^[99]^ Next, analyte molecules were filtered and injected at a flow rate of 30-50 µL/min over a decorin- or collagen II-immobilized surface at a wide-range of descending molar concentrations at 3× dilution, including 1,200-1.646 nM for collagen II, decorin, aggrecan core protein, and 400-0.549 nM for intact aggrecan. To re-establish baseline signals between analyte injections, the surface was regenerated with two repeats of PBS-P buffer (blank) injections and 12-sec + 7-sec pulses of glycine (pH = 2) between different concentrations for the same analyte, and four repeats when switching to different analytes. Each binding assay was repeated in triplicate on independent immobilization cycles to minimize surface-specific variability.^[100]^

Raw sensorgrams were recorded at 40 Hz to capture fast-binding events,^[101]^ and post-processed in Scrubber 2.0c (BioLogic Software)^[100]^ through reference channel and blank subtraction, baseline alignment and normalization across analyte concentrations to minimize instrumental noise and sample refractive index variability. For each ligand-analyte pair, the processed sensorgrams from multiple replicates (*n* = 3) were fitted in ClampXP (BioLogic)^[102]^ through Monte Carlo simulations (≤ 50 iterations), resampling sensorgrams within experimental noise envelope^[103]^ and refitting each iteration to evaluate the goodness-of-fit of relevant binding interaction models^[104]^ (Table 2^[49,52–53,56]^). For each ligand-analyte pair, selection of the appropriate kinetics model (Table 2) was based on the model yielding the minimal residual errors denoted by χ^2^-values representing noise-weighted least squares deviation between experimental and simulated sensorgrams in ClampXP.^[102,105]^ Based on the selected model (Table 3), the predicted kinetic parameters, association rate (*k_a_*), dissociation rate (*k_d_*) and equilibrium dissociation constant (*K_D_*) were calculated for each best-fitted model via the Monte Carlo simulation (Table 3).

The three-component “ternary” SPR setup was applied to assess whether decorin acts as a coupler to stabilize collagen II-aggrecan interactions. Following the same procedure as for the binary setup, we immobilized collagen II or intact aggrecan as the ligand molecule onto the PEG-based sensor at pH 7.4 at a concentration of 100-200 µg/mL in 1× PBS via EDC-NHS chemistry. Collagen II or aggrecan analytes were first injected at descending concentrations, with surface regenerated between injections. Next, the same series of analyte injections were repeated, but with each injection preceded by the injection of 400 nM decorin solution to enable ligand-decorin binding as the coupler (Fig. 5d). The concentration of decorin used was the highest concentration that yielded stable bindings with collagen II and aggrecan measured from the binary setup.

Due to the inherent complex nature of ternary binding interactions, we did not apply Monte Carlo simulations to multiple sensorgrams to derive the binding kinetics. Instead, after following the same procedure for instrumental noise minimization as for the binary setup, we applied the classic 1:1 Langmuir model^[56]^ to individual sets of sensorgrams with Scrubber using a dissociation-weighted fitting approach^[100]^ to compare the fold changes in apparent *K_D_* for ligand-analyte binding with versus without decorin as the coupler (Table S10). These apparent *K_D_*’s derived from Scrubber were different from predicted *K_D_*’s derived the Monte Carlo simulation on multiple replicates of sensorgrams. Also, the observed large variations in apparent *K_D_*’s across multiple replicates (Table S10) were expected due to variations in the heterogeneity of surface ligand density and analyte mass transport.^[49]^ However, since fold changes in apparent *K_D_*’s were calculated within each individual set, effects of decorin on relative aggrecan-collagen binding affinities were not obscured by sample-to-sample variations.

### AFM-based nanoindentation

Freshly dissected femoral condyle cartilage from 3-week-old WT and *Dcn^−/−^* mice were digested with a cocktail of 0.5 U/mL chondroitinase ABC and 500 U/mL hyaluronidase in 1× PBS for 24 hours to remove GAGs. Either untreated or GAG-depleted condyles were incubated with 20 µg/mL bovine decorin in 1× PBS for 24 hours to allow for the infiltration of exogenous decorin. AFM-nanoindentation was performed on untreated and GAG-depleted samples, with or without the infiltration of decorin (Fig. 6c, *n* ≥ 10). In a separate group, additional condyles were stimulated by 10 ng/mL IL-1β for 24 hrs, followed by GAG removal and GAG removal with decorin infiltration, and then, subjected to AFM-nanoindentation (Fig. 6d, *n* ≥ 10). Indentation tests were performed on a Dimension Icon AFM (BrukerNano) using borosilicate microspherical colloidal tips (*R* ≈ 5 μm, nominal *k* ≈ 8.9 N/m, HQ:NSC35/tipless/Cr-Au, cantilever A, NanoAndMore) at 10 μm/s rate up to ≈ 1 μN maximum load in 1× PBS with protease inhibitors (Pierce A32965, ThermoFisher), following the established procedure.^[30]^ For each sample, at least 10-15 randomly chosen locations were tested on the load-bearing regions of the medial condyle to account for spatial heterogeneity. The effective indentation modulus, *E*_ind_, was calculated by fitting the entire loading portion of each force-indentation depth (*F-D*) curve to the Hertz model.^[106–107]^

### Statistical analysis

The linear mixed model was applied to analyze spatial parameters, *I*_mean_, *I*_peak_, *d*_peak_, *d*_1/2_ and *d*_peak_ + *d*_1/2_, as well as the cumulative release of nascent GAGs or proteins using the R package lme4 (version 1.1-35).^[108]^ In these tests, genotype (WT versus *Dcn^−/−^*), stimulation condition (IL-1β versus untreated), decorin pre/post-treatment groups were treated as between-subject fixed-effect factors when appropriate, and time point (day 0 to day 6 or 8) was treated as a within-subject fixed-effect factor, with interaction terms between the fixed effects. For spatial parameters, given the analysis of multiple cells per animal, individual animal was treated as a random-effect factor. To account for family-wise type I errors, Holm-Bonferroni multiple contrast correction was applied to calculate the adjusted *p*-values between genotypes and IL-1β stimulation conditions, and Tukey-Kramer post-hoc comparison was applied to calculate adjusted *p*-values among pre/post-treatments of decorin and across all time points for each genotype, IL-1β stimulation or treatment condition. For indentation modulus, *E*_ind_, the linear mixed model was applied to test the effects of treatments (GAG-depletion and decorin infiltration), genotype and IL-1β stimulation. Similarly, Tukey-Kramer comparison was applied to calculate the adjusted *p*-values between treatments, and Holm-Bonferroni correction was applied between genotype and IL-1β stimulation. In all the tests, the significance level was set at *α* = 0.05. A complete list of all quantitative and statistical outcomes is summarized in Table 3 and Tables S1–S11.

## Acknowledgements

This work was financially supported by the National Institutes of Health (NIH) Grant R01AR074490 (to LH), R01AR074472 (to XLL), as well as NIH Grant P30AR069619 to the Penn Center for Musculoskeletal Disorders (PCMD). We thank the Cell Imaging Center at Drexel University for the use of the Zeiss LSM700 confocal microscope.

## References

[1] GBD 2021 Osteoarthritis Collaborators, Global, regional, and national burden of osteoarthritis, 1990-2020 and projections to 2050: a systematic analysis for the Global Burden of Disease Study 2021, Lancet Rheumatol 2023, 5, e508.

[2] Y. Krishnan, A. J. Grodzinsky, Cartilage diseases, Matrix Biol. 2018, 71-72, 51.

[3] A. Maroudas, in (Ed.: M. A. R. Freeman), Pitman, England 1979.

[4] L. Han, A. J. Grodzinsky, C. Ortiz, Nanomechanics of the cartilage extracellular matrix, Annu. Rev. Mater. Res. 2011, 41, 133.

[5] F. Guilak, R. J. Nims, A. Dicks, C. L. Wu, I. Meulenbelt, Osteoarthritis as a disease of the cartilage pericellular matrix, Matrix Biol. 2018, 71-72, 40.

[6] T. E. Hardingham, H. Muir, The specific interaction of hyaluronic acid with cartilage proteoglycans, Biochim. Biophys. Acta 1972, 279, 401.

[7] J. A. Buckwalter, L. C. Rosenberg, L.-H. Tang, The effect of link protein on proteoglycan aggregate structure - an electron microscopic study of the molecular architecture and dimensions of proteoglycan aggregates reassembled from the proteoglycan monomers and link proteins of bovine fetal epiphyseal cartilage, J. Biol. Chem. 1984, 259, 5361.

[8] T. N. Wight, D. K. Heinegård, V. C. Hascall, Cell Biology of Extracellular Matrix, Plenum press, New York 1991.

[9] V. C. Mow, S. C. Kuei, W. M. Lai, C. G. Armstrong, Biphasic creep and stress relaxation of articular cartilage in compression: theory and experiments, J. Biomech. Eng. 1980, 102, 73.

[10] A. K. Williamson, A. C. Chen, R. L. Sah, Compressive properties and function-composition relationships of developing bovine articular cartilage, J. Orthop. Res. 2001, 19, 1113.

[11] W. B. Zhu, J. C. Iatridis, V. Hlibczuk, A. Ratcliffe, V. C. Mow, Determination of collagen-proteoglycan interactions in vitro, J. Biomech. 1996, 29, 773.

[12] T. Wyse Jackson, J. Michel, P. Lwin, L. A. Fortier, M. Das, L. J. Bonassar, I. Cohen, Structural origins of cartilage shear mechanics, Sci Adv 2022, 8, eabk2805.

[13] L. S. Lohmander, M. Ionescu, H. Jugessur, A. R. Poole, Changes in joint cartilage aggrecan after knee injury and in osteoarthritis, Arthritis Rheum 1999, 42, 534.

[14] M. T. Bayliss, S. Howat, C. Davidson, J. Dudhia, The organization of aggrecan in human articular cartilage. Evidence for age-related changes in the rate of aggregation of newly synthesized molecules., J. Biol. Chem. 2000, 275, 6321.

[15] T. M. Quinn, A. A. Maung, A. J. Grodzinsky, E. B. Hunziker, J. D. Sandy, Physical and biological regulation of proteoglycan turnover around chondrocytes in cartilage explants. Implications for tissue degradation and repair, Ann. N. Y. Acad. Sci. 1999, 878, 420.

[16] N. Verzijl, J. DeGroot, S. R. Thorpe, R. A. Bank, J. N. Shaw, T. J. Lyons, J. W. J. Bijlsma, F. P. J. G. Lafeber, J. W. Baynes, J. M. TeKoppele, Effect of collagen turnover on the accumulation of advanced glycation end products, J. Biol. Chem. 2000, 275, 39027.

[17] M. W. Lark, J. T. Gordy, J. R. Weidner, J. Ayala, J. H. Kimura, H. R. Williams, R. A. Mumford, C. R. Flannery, S. S. Carlson, M. Iwata, J. D. Sandy, Cell-mediated catabolism of aggrecan evidence that cleavage at the aggrecanase site (Glu^373^-Ala^374^) proteolysis of the interglobular domain, J. Biol. Chem. 1995, 270, 2550.

[18] I. Singer, D. W. Kawka, E. K. Bayne, S. A. Donatelli, J. R. Weidner, H. R. Williams, J. M. Ayala, R. A. Mumford, M. W. Lark, T. T. Glant, VDIPEN, a metalloproteinase-generated neoepitope, is induced and immunolocalized in articular cartilage during inflammatory arthritis, J. Clin. Invest. 1995, 95, 2178.

[19] I. G. Otterness, J. T. Downs, C. Lane, M. L. Bliven, H. Stukenbrok, D. N. Scampoli, A. J. Milici, P. S. Mezes, Detection of collagenase-induced damage of collagen by 9A4, a monoclonal C-terminal neoepitope antibody, Matrix Biol. 1999, 18, 331.

[20] P. J. Roughley, J. S. Mort, The role of aggrecan in normal and osteoarthritic cartilage, J. Exp. Orthop. 2014, 1, 8.

[21] D. R. Chery, B. Han, Q. Li, Y. Zhou, S. J. Heo, B. Kwok, P. Chandrasekaran, C. Wang, L. Qin, X. L. Lu, D. Kong, M. Enomoto-Iwamoto, R. L. Mauck, L. Han, Early changes in cartilage pericellular matrix micromechanobiology portend the onset of post-traumatic osteoarthritis, Acta Biomater. 2020, 111, 267.

[22] M. B. Goldring, S. R. Goldring, Articular cartilage and subchondral bone in the pathogenesis of osteoarthritis, Ann. N. Y. Acad. Sci. 2010, 1192, 230.

[23] Z. Peng, H. Sun, V. Bunpetch, Y. Koh, Y. Wen, D. Wu, H. Ouyang, The regulation of cartilage extracellular matrix homeostasis in joint cartilage degeneration and regeneration, Biomaterials 2021, 268, 120555.

[24] A. R. Poole, L. C. Rosenberg, A. Reiner, M. Ionescu, E. Bogoch, P. J. Roughley, Contents and distributions of the proteoglycans decorin and biglycan in normal and osteoarthritic human articular cartilage, J. Orthop. Res. 1996, 14, 681.

[25] A. McAlinden, J. Dudhia, M. C. Bolton, P. Lorenzo, D. Heinegård, M. T. Bayliss, Age-related changes in the synthesis and mRNA expression of decorin and aggrecan in human meniscus and articular cartilage, Osteoarthritis Cartilage 2001, 9, 33.

[26] O. Sampaio Lde, M. T. Bayliss, T. E. Hardingham, H. Muir, Dermatan sulphate proteoglycan from human articular cartilage. Variation in its content with age and its structural comparison with a small chondroitin sulphate proteoglycan from pig laryngeal cartilage, Biochem J 1988, 254, 757.

[27] G. Cs-Szabó, P. J. Roughley, A. H. K. Plaas, T. T. Glant, Large and small proteoglycans of osteoarthritic and rheumatoid articular-cartilage, Arthritis Rheum. 1995, 38, 660.

[28] G. Cs-Szabó, L. I. Melching, P. J. Roughley, T. T. Glant, Changes in messenger RNA and protein levels of proteoglycans and link protein in human osteoarthritic cartilage samples, Arthritis Rheum. 1997, 40, 1037.

[29] G. Barreto, A. Soininen, P. Ylinen, J. Sandelin, Y. T. Konttinen, D. C. Nordstrom, K. K. Eklund, Soluble biglycan: a potential mediator of cartilage degradation in osteoarthritis, Arthritis Res Ther 2015, 17, 379.

[30] B. Han, Q. Li, C. Wang, P. Patel, S. M. Adams, B. Doyran, H. T. Nia, R. Oftadeh, S. Zhou, C. Y. Li, X. S. Liu, X. L. Lu, M. Enomoto-Iwamoto, L. Qin, R. L. Mauck, R. V. Iozzo, D. E. Birk, L. Han, Decorin regulates the aggrecan network integrity and biomechanical functions of cartilage extracellular matrix, ACS Nano 2019, 13, 11320.

[31] D. R. Chery, B. Han, Y. Zhou, C. Wang, S. M. Adams, P. Chandrasekaran, B. Kwok, S.-J. Heo, M. Enomoto-Iwamoto, X. L. Lu, D. Kong, R. V. Iozzo, D. E. Birk, R. L. Mauck, L. Han, Decorin regulates cartilage pericellular matrix micromechanobiology, Matrix Biol. 2021, 96, 1.

[32] S. S. Glasson, T. J. Blanchet, E. A. Morris, The surgical destabilization of the medial meniscus (DMM) model of osteoarthritis in the 129/SvEv mouse, Osteoarthritis Cartilage 2007, 15, 1061.

[33] Q. Li, B. Han, C. Wang, W. Tong, W. J. Tseng, L.-H. Han, X. S. Liu, M. Enomoto-Iwamoto, R. L. Mauck, L. Qin, R. V. Iozzo, D. E. Birk, L. Han, Mediation of cartilage matrix degeneration and fibrillation by decorin in post-traumatic osteoarthritis, Arthritis Rheumatol. 2020, 72, 1266.

[34] B. Han, Q. Li, C. Wang, P. Chandrasekaran, Y. Zhou, L. Qin, X. S. Liu, M. Enomoto-Iwamoto, D. Kong, R. V. Iozzo, D. E. Birk, L. Han, Differentiated activities of decorin and biglycan in the progression of post-traumatic osteoarthritis, Osteoarthritis Cartilage 2021, 29, 1181.

[35] D. R. Keene, J. D. San Antonio, R. Mayne, D. J. McQuillan, G. Sarris, S. A. Santoro, R. V. Iozzo, Decorin binds near the C terminus of type I collagen, J Biol Chem 2000, 275, 21801.

[36] M. A. Gubbiotti, S. D. Vallet, S. Ricard-Blum, R. V. Iozzo, Decorin interacting network: A comprehensive analysis of decorin-binding partners and their versatile functions, Matrix Biol. 2016, 55, 7.

[37] A. Hildebrand, M. Romarís, L. M. Rasmussen, D. Heinegård, D. R. Twardzik, W. A. Border, E. Ruoslahti, Interaction of the small interstitial proteoglycans biglycan, decorin and fibromodulin with transforming growth factor *β*, Biochem. J. 1994, 302, 527.

[38] C. Loebel, R. L. Mauck, J. A. Burdick, Local nascent protein deposition and remodelling guide mesenchymal stromal cell mechanosensing and fate in three-dimensional hydrogels, Nat. Mater. 2019, 18, 883.

[39] C. Loebel, M. Y. Kwon, C. Wang, L. Han, R. L. Mauck, J. A. Burdick, Metabolic labeling to probe the spatiotemporal accumulation of matrix at the chondrocyte-hydrogel interface, Adv. Funct. Mater. 2020, 30, 1909802.

[40] A. Porter, L. Wang, L. Han, X. L. Lu, Bio-orthogonal click chemistry methods to evaluate the metabolism of inflammatory challenged cartilage after traumatic overloading, ACS Biomater. Sci. Eng. 2022, 8, 2564.

[41] K. G. Danielson, H. Baribault, D. F. Holmes, H. Graham, K. E. Kadler, R. V. Iozzo, Targeted disruption of decorin leads to abnormal collagen fibril morphology and skin fragility, J. Cell Biol. 1997, 136, 729.

[42] T. Gronau, K. Kruger, C. Prein, A. Aszodi, I. Gronau, R. V. Iozzo, F. C. Mooren, H. Clausen-Schaumann, J. Bertrand, T. Pap, P. Bruckner, R. Dreier, Forced exercise-induced osteoarthritis is attenuated in mice lacking the small leucine-rich proteoglycan decorin, Ann. Rheum. Dis. 2017, 76, 442.

[43] K. W. Finnson, Y. Almadani, A. Philip, Non-canonical (non-SMAD2/3) TGF-β signaling in fibrosis: Mechanisms and targets, Semin Cell Dev Biol 2020, 101, 115.

[44] M. Santra, C. C. Reed, R. V. Iozzo, Decorin binds to a narrow region of the epidermal growth factor (EGF) receptor, partially overlapping but distinct from the EGF-binding epitope, J. Biol. Chem. 2002, 277, 35671.

[45] N. Lala, G. V. Girish, A. Cloutier-Bosworth, P. K. Lala, Mechanisms in decorin regulation of vascular endothelial growth factor-induced human trophoblast migration and acquisition of endothelial phenotype, Biol Reprod 2012, 87, 59.

[46] S. Goldoni, A. Humphries, A. Nystrom, S. Sattar, R. T. Owens, D. J. McQuillan, K. Ireton, R. V. Iozzo, Decorin is a novel antagonistic ligand of the Met receptor, J. Cell Biol. 2009, 185, 743.

[47] L. Li, M. Ly, R. J. Linhardt, Proteoglycan sequence, Mol Biosyst 2012, 8, 1613.

[48] J. Carroll, M. Raum, K. Forsten-Williams, U. C. Täuber, Ligand-receptor binding kinetics in surface plasmon resonance cells: a Monte Carlo analysis, Phys Biol 2016, 13, 066010.

[49] J. Svitel, H. Boukari, D. Van Ryk, R. C. Willson, P. Schuck, Probing the functional heterogeneity of surface binding sites by analysis of experimental binding traces and the effect of mass transport limitation, Biophys J 2007, 92, 1742.

[50] R. Tenni, M. Viola, F. Welser, P. Sini, C. Giudici, A. Rossi, M. E. Tira, Interaction of decorin with CNBr peptides from collagens I and II. Evidence for multiple binding sites and essential lysyl residues in collagen, Eur J Biochem 2002, 269, 1428.

[51] P. G. Scott, P. A. McEwan, C. M. Dodd, E. M. Bergmann, P. N. Bishop, J. Bella, Crystal structure of the dimeric protein core of decorin, the archetypal small leucine-rich repeat proteoglycan, Proc. Natl. Acad. Sci. USA 2004, 101, 15633.

[52] T. A. Morton, D. G. Myszka, Kinetic analysis of macromolecular interactions using surface plasmon resonance biosensors, Methods Enzymol 1998, 295, 268.

[53] K. M. Müller, K. M. Arndt, A. Plückthun, Model and simulation of multivalent binding to fixed ligands, Anal. Biochem. 1998, 261, 149.

[54] H. Watanabe, S. C. Cheung, N. Itano, K. Kimata, Y. Yamada, Identification of hyaluronan-binding domains of aggrecan, J. Biol. Chem. 1997, 272, 28057.

[55] A. Aspberg, S. Adam, G. Kostka, R. Timpl, D. Heinegård, Fibulin-1 is a ligand for the C-type lectin domains of aggrecan and versican, J Biol Chem 1999, 274, 20444.

[56] I. Langmuir, The constitution and fundamental properties of solids and liquids. Part I. Solids, J. Am. Chem. Soc. 1916, 38, 2221.

[57] M. J. Roy, S. Winkler, S. J. Hughes, C. Whitworth, M. Galant, W. Farnaby, K. Rumpel, A. Ciulli, SPR-measured dissociation kinetics of PROTAC ternary complexes influence target degradation rate, ACS Chemical Biology 2019, 14, 361.

[58] L. Han, E. H. Frank, J. J. Greene, H.-Y. Lee, H.-H. K. Hung, A. J. Grodzinsky, C. Ortiz, Time-dependent nanomechanics of cartilage, Biophys. J. 2011, 100, 1846.

[59] H.-Y. Lee, L. Han, P. J. Roughley, A. J. Grodzinsky, C. Ortiz, Age-related nanostructural and nanomechanical changes of individual human cartilage aggrecan monomers and their glycosaminoglycan side chains, J. Struct. Biol. 2013, 181, 264.

[60] M. W. Lark, E. K. Bayne, J. Flanagan, C. F. Harper, L. A. Hoerrner, N. I. Hutchinson, Singer, II, S. A. Donatelli, J. R. Weidner, H. R. Williams, R. A. Mumford, L. S. Lohmander, Aggrecan degradation in human cartilage. Evidence for both matrix metalloproteinase and aggrecanase activity in normal, osteoarthritic, and rheumatoid joints, J. Clin. Invest. 1997, 100, 93.

[61] T. Yasumoto, J. L. E. Bird, K. Sugimoto, R. M. Mason, M. T. Bayliss, The G1 domain of aggrecan released from porcine articular cartilage forms stable complexes with hyaluronan/link protein, Rheumatology 2003, 42, 336.

[62] S. Chen, D. E. Birk, The regulatory roles of small leucine-rich proteoglycans in extracellular matrix assembly, FEBS J. 2013, 280, 2120.

[63] D. R. Eyre, M. A. Weis, J.-J. Wu, Articular cartilage collagen: an irreplaceable framework?, Eur. Cell. Mater. 2006, 12, 57.

[64] C. Wang, B. K. Brisson, M. Terajima, Q. Li, K. Hoxha, B. Han, A. M. Goldberg, X. Sherry Liu, M. S. Marcolongo, M. Enomoto-Iwamoto, M. Yamauchi, S. W. Volk, L. Han, Type III collagen is a key regulator of the collagen fibrillar structure and biomechanics of articular cartilage and meniscus, Matrix Biol. 2020, 85-86, 47.

[65] L. Han, D. Dean, L. A. Daher, A. J. Grodzinsky, C. Ortiz, Cartilage aggrecan can undergo self-adhesion, Biophys. J. 2008, 95, 4862.

[66] F. P. Rojas, M. A. Batista, C. A. Lindburg, D. Dean, A. J. Grodzinsky, C. Ortiz, L. Han, Molecular adhesion between cartilage extracellular matrix macromolecules, Biomacromolecules 2014, 15, 772.

[67] M. Islam, J. Gor, S. J. Perkins, Y. Ishikawa, H. P. Bächinger, E. Hohenester, The concave face of decorin mediates reversible dimerization and collagen binding, J Biol Chem 2013, 288, 35526.

[68] L. Ng, A. J. Grodzinsky, P. Patwari, J. Sandy, A. Plaas, C. Ortiz, Individual cartilage aggrecan macromolecules and their constituent glycosaminoglycans visualized via atomic force microscopy, J. Struct. Biol. 2003, 143, 242.

[69] A. Aruffo, I. Stamenkovic, M. Melnick, C. B. Underhill, B. Seed, CD44 is the principal cell surface receptor for hyaluronate, Cell 1990, 61, 1303.

[70] R. E. Wilusz, J. Sanchez-Adams, F. Guilak, The structure and function of the pericellular matrix of articular cartilage, Matrix Biol. 2014, 39, 25.

[71] R. J. Wenstrup, J. B. Florer, E. W. Brunskill, S. M. Bell, I. Chervoneva, D. E. Birk, Type V collagen controls the initiation of collagen fibril assembly, J. Biol. Chem. 2004, 279, 53331.

[72] C. Wang, M. Fan, S. C. Heo, S. M. Adams, T. Li, Y. Liu, Q. Li, C. Loebel, J. A. Burdick, X. L. Lu, D. E. Birk, F. Alisafaei, R. L. Mauck, L. Han, Structure, mechanics, and mechanobiology of fibrocartilage pericellular matrix mediated by type V collagen, Adv. Sci. 2025, 12, e14750.

[73] F. W. Roemer, D. T. Felson, J. J. Stefanik, G. Rabasa, N. Wang, M. D. Crema, T. Neogi, M. C. Nevitt, J. Torner, C. E. Lewis, C. Peloquin, A. Guermazi, Heterogeneity of cartilage damage in Kellgren and Lawrence grade 2 and 3 knees: the MOST study, Osteoarthritis Cartilage 2022, 30, 714.

[74] E. R. Kahle, B. Han, P. Chandrasekaran, E. R. Phillips, M. K. Mulcahey, X. L. Lu, M. S. Marcolongo, L. Han, Molecular engineering of pericellular microniche via biomimetic proteoglycans modulates cell mechanobiology, ACS Nano 2022, 16, 1220.

[75] S. G. Lopez, H. R. Moura, E. Chow, J. C.-H. Kuo, M. J. Paszek, L. J. Bonassar, Recombinant small leucine-rich proteoglycans modulate fiber structure and mechanical properties of collagen gels, ACS Biomaterials Science & Engineering 2025.

[76] H. I. Ma, D. Y. Hueng, H. A. Shui, J. M. Han, C. H. Wang, Y. H. Lai, S. Y. Cheng, X. Xiao, M. T. Chen, Y. P. Yang, Intratumoral decorin gene delivery by AAV vector inhibits brain glioblastomas and prolongs survival of animals by inducing cell differentiation, Int J Mol Sci 2014, 15, 4393.

[77] S. S. Sivan, E. Wachtel, P. Roughley, Structure, function, aging and turnover of aggrecan in the intervertebral disc, Biochim Biophys Acta 2014, 1840, 3181.

[78] K. Yoshida, S. Takatsuka, E. Hatada, H. Nakamura, A. Tanaka, K. Ueki, K. Nakagawa, Y. Okada, E. Yamamoto, R. Fukuda, Expression of matrix metalloproteinases and aggrecanase in the synovial fluids of patients with symptomatic temporomandibular disorders, Oral Surg Oral Med Oral Pathol Oral Radiol Endod 2006, 102, 22.

[79] H. Port, B. Coppers, S. Tragl, E. Manger, L. M. Niemiec, S. Bayat, D. Simon, F. Fagni, G. Corte, A. C. Bay-Jensen, K. Tascilar, A. J. Hueber, K. G. Schmidt, V. Schönau, M. Sticherling, S. Heinrich, S. Leyendecker, D. Bohr, G. Schett, A. Kleyer, S. Holm Nielsen, A. M. Liphardt, Serum extracellular matrix biomarkers in rheumatoid arthritis, psoriatic arthritis and psoriasis and their association with hand function, Sci Rep 2025, 15, 13656.

[80] B. G. Gibson, M. D. Briggs, The aggrecanopathies; an evolving phenotypic spectrum of human genetic skeletal diseases, Orphanet J Rare Dis 2016, 11, 86.

[81] I. T. Weber, R. W. Harrison, R. V. Iozzo, Model structure of decorin and implications for collagen fibrillogenesis, J Biol Chem 1996, 271, 31767.

[82] K. Baghy, H. Szakadáti, I. Kovalszky, Decorin the antifibrotic proteoglycan and its progression in therapy, Am J Physiol Cell Physiol 2025, 328, C1853.

[83] H. Järveläinen, P. Puolakkainen, S. Pakkanen, E. L. Brown, M. Höök, R. V. Iozzo, E. H. Sage, T. N. Wight, A role for decorin in cutaneous wound healing and angiogenesis, Wound Repair Regen 2006, 14, 443.

[84] D. D. Sofeu Feugaing, M. Götte, M. Viola, More than matrix: the multifaceted role of decorin in cancer, Eur J Cell Biol 2013, 92, 1.

[85] D. Kim, B. Langmead, S. L. Salzberg, HISAT: a fast spliced aligner with low memory requirements, Nat Methods 2015, 12, 357.

[86] M. Pertea, G. M. Pertea, C. M. Antonescu, T. C. Chang, J. T. Mendell, S. L. Salzberg, StringTie enables improved reconstruction of a transcriptome from RNA-seq reads, Nat Biotechnol 2015, 33, 290.

[87] C. Trapnell, A. Roberts, L. Goff, G. Pertea, D. Kim, D. R. Kelley, H. Pimentel, S. L. Salzberg, J. L. Rinn, L. Pachter, Differential gene and transcript expression analysis of RNA-seq experiments with TopHat and Cufflinks, Nat Protoc 2012, 7, 562.

[88] L. Kong, Y. Zhang, Z. Q. Ye, X. Q. Liu, S. Q. Zhao, L. Wei, G. Gao, CPC: assess the protein-coding potential of transcripts using sequence features and support vector machine, Nucleic Acids Res 2007, 35, W345.

[89] A. McKenna, M. Hanna, E. Banks, A. Sivachenko, K. Cibulskis, A. Kernytsky, K. Garimella, D. Altshuler, S. Gabriel, M. Daly, M. A. DePristo, The Genome Analysis Toolkit: a MapReduce framework for analyzing next-generation DNA sequencing data, Genome Res 2010, 20, 1297.

[90] B. Langmead, S. L. Salzberg, Fast gapped-read alignment with Bowtie 2, Nat Methods 2012, 9, 357.

[91] M. I. Love, W. Huber, S. Anders, Moderated estimation of fold change and dispersion for RNA-seq data with DESeq2, Genome Biol 2014, 15, 550.

[92] R. O. Hynes, A. Naba, Overview of the matrisome—an inventory of extracellular matrix constituents and functions, Cold Spring Harb Perspect Biol 2012, 4, a004903.

[93] D. Tang, M. Chen, X. Huang, G. Zhang, L. Zeng, G. Zhang, S. Wu, Y. Wang, SRplot: A free online platform for data visualization and graphing, PLoS One 2023, 18, e0294236.

[94] H. Stanton, S. B. Golub, F. M. Rogerson, K. Last, C. B. Little, A. J. Fosang, Investigating ADAMTS-mediated aggrecanolysis in mouse cartilage, Nat. Protoc. 2011, 6, 388.

[95] G. A. Canziani, J. A. Melero, E. R. Lacy, Characterization of neutralizing affinity-matured human respiratory syncytial virus F binding antibodies in the sub-picomolar affinity range, J Mol Recognit 2012, 25, 136.

[96] S. Zhang, A. P. Holmes, A. Dick, A. A. Rashad, L. Enríquez Rodríguez, G. A. Canziani, M. J. Root, I. M. Chaiken, Altered Env conformational dynamics as a mechanism of resistance to peptide-triazole HIV-1 inactivators, Retrovirology 2021, 18, 31.

[97] A. Nangarlia, F. F. Hassen, G. Canziani, P. Bandi, C. Talukder, F. Zhang, D. Krauth, E. N. Gary, D. B. Weiner, P. Bieniasz, S. Navas-Martin, B. R. O’Keefe, C. G. Ang, I. Chaiken, Irreversible Inactivation of SARS-CoV-2 by Lectin Engagement with Two Glycan Clusters on the Spike Protein, Biochemistry 2023, 62, 2115.

[98] K. Uchida, H. Otsuka, M. Kaneko, K. Kataoka, Y. Nagasaki, A reactive poly(ethylene glycol) layer to achieve specific surface plasmon resonance sensing with a high s/n ratio: The substantial role of a short underbrushed PEG layer in minimizing nonspecific adsorption, Analytical Chemistry 2005, 77, 1075.

[99] G. Galicia-Vázquez, L. Lindqvist, X. Wang, I. Harvey, J. Liu, J. Pelletier, High-throughput assays probing protein-RNA interactions of eukaryotic translation initiation factors, Anal Biochem 2009, 384, 180.

[100] D. G. Myszka, Improving biosensor analysis, J. Mol. Recognit. 1999, 12, 279.

[101] R. Karlsson, V. Fridh, Å. Frostell, Surrogate potency assays: Comparison of binding profiles complements dose response curves for unambiguous assessment of relative potencies, J Pharm Anal 2018, 8, 138.

[102] D. G. Myszka, T. A. Morton, CLAMP: a biosensor kinetic data analysis program, Trends Biochem Sci 1998, 23, 149.

[103] R. Karlsson, A. Fält, Experimental design for kinetic analysis of protein-protein interactions with surface plasmon resonance biosensors, J Immunol Methods 1997, 200, 121.

[104] J. Svitel, A. Balbo, R. A. Mariuzza, N. R. Gonzales, P. Schuck, Combined affinity and rate constant distributions of ligand populations from experimental surface binding kinetics and equilibria, Biophys J 2003, 84, 4062.

[105] P. Schuck, Use of surface plasmon resonance to probe the equilibrium and dynamic aspects of interactions between biological macromolecules, Annu Rev Biophys Biomol Struct 1997, 26, 541.

[106] B. Doyran, W. Tong, Q. Li, H. Jia, X. Zhang, C. Chen, M. Enomoto-Iwamoto, X. L. Lu, L. Qin, L. Han, Nanoindentation modulus of murine cartilage: a sensitive indicator of the initiation and progression of post-traumatic osteoarthritis, Osteoarthritis Cartilage 2017, 25, 108.

[107] B. Han, H. T. Nia, C. Wang, P. Chandrasekaran, Q. Li, D. R. Chery, H. Li, A. J. Grodzinsky, L. Han, AFM-nanomechanical test: an interdisciplinary tool that links the understanding of cartilage and meniscus biomechanics, osteoarthritis degeneration and tissue engineering, ACS Biomater. Sci. Eng. 2017, 3, 2033.

[108] D. Bates, M. Mächler, B. Bolker, S. Walker, Fitting linear mixed-effects models using lme4, J. Stat. Softw. 2015, 67(1), 1.

