## Supporting Materials for "Decorin Promotes Nascent Proteoglycan Retention in Cartilage Matrix by Strengthening Collagen II-Aggrecan Integration"

Dr. Lin Han

.

**Table S1.** Summary of averaged values and statistical analysis outcomes of the mean GAL intensity, $I_{\text{mean}}$, extracted from high-resolution confocal images of click-labeled femoral head cartilage explants of age-matched wild-type (WT) and decorin-null ($Dcn^{-/-}$) mice over 6 days of culture, shown as mean ± 95% CI from values averaged by each explant. The $p$-values were calculated using the linear mixed-effect model, followed by Holm-Bonferroni correction for multiple contrasts between genotypes and treatments, and Tukey-Kramer correction for multiple comparisons across culture time points for each genotype and treatment.

| $I_{\text{mean}}$ (GAL) (a.u.) | Untreated | | | | | |
|---|---|---|---|---|---|---|
| | WT | | | $Dcn^{-/-}$ | | |
| | mean ± 95% CI | *n* (cells) | *N* (animals) | mean ± 95% CI | *n* (cells) | *N* (animals) |
| Day 0 | 39 ± 2 | 114 | 4 | 36 ± 1 | 95 | 4 |
| Day 2 | 41 ± 2 | 78 | 4 | 31 ± 1 | 110 | 4 |
| Day 4 | 39 ± 8 | 111 | 4 | 31 ± 1 | 108 | 4 |
| Day 6 | 34 ± 1 | 104 | 4 | 26 ± 1 | 102 | 4 |
| $I_{\text{mean}}$ (GAL) (a.u.) | IL-1β | | | | | |
| | WT | | | $Dcn^{-/-}$ | | |
| | mean ± 95% CI | *n* (cells) | *N* (animals) | mean ± 95% CI | *n* (cells) | *N* (animals) |
| Day 2 | 29 ± 1 | 131 | 4 | 27 ± 1 | 111 | 4 |
| Day 4 | 23 ± 1 | 114 | 4 | 22 ± 1 | 109 | 4 |
| Day 6 | 20 ± 1 | 91 | 4 | 19 ± 1 | 97 | 4 |

| *p*-value (genotype/IL-1β) | WT vs $Dcn^{-/-}$ | | Untreated vs IL-1β | |
|---|---|---|---|---|
| | Control | IL-1β | WT | $Dcn^{-/-}$ |
| Day 0 | 0.320 | -- | -- | -- |
| Day 2 | 0.049 | 0.656 | 0.030 | 0.313 |
| Day 4 | 0.034 | 0.825 | < 0.001 | 0.017 |
| Day 6 | 0.038 | 0.760 | 0.010 | 0.011 |

| *p*-value (time points) | Untreated | | IL-1β | |
|---|---|---|---|---|
| | WT | $Dcn^{-/-}$ | WT | $Dcn^{-/-}$ |
| Day 0 vs 2 | 0.990 | 0.741 | -- | -- |
| Day 0 vs 4 | 0.958 | 0.585 | -- | -- |
| Day 0 vs 6 | 0.217 | 0.073 | -- | -- |
| Day 2 vs 4 | 0.852 | 0.993 | 0.139 | 0.271 |
| Day 2 vs 6 | 0.137 | 0.350 | 0.048 | 0.084 |
| Day 4 vs 6 | 0.428 | 0.486 | 0.770 | 0.714 |

**Table S2.** Summary of averaged values and statistical analysis outcomes of the mean AHA intensity, $I_{mean}$, extracted from high-resolution confocal images of click-labeled femoral head cartilage explants of age-matched wild-type (WT) and decorin-null (*Dcn*$^{-/-}$) mice over 6 days of culture, shown as mean ± 95% CI from values averaged by each explant. The *p*-values were calculated using the linear mixed-effect model, followed by Holm-Bonferroni correction for multiple contrasts between genotypes and treatments, and Tukey-Kramer correction for multiple comparisons across culture time points for each genotype and treatment.

| $I_{mean}$ (AHA) (a.u.) | Untreated | | | | | |
|---|---|---|---|---|---|---|
| | WT | | | *Dcn*$^{-/-}$ | | |
| | mean ± 95% CI | *n* (cells) | *N* (animals) | mean ± 95% CI | *n* (cells) | *N* (animals) |
| Day 0 | 34 ± 2 | 76 | 3 | 35 ± 2 | 115 | 3 |
| Day 2 | 34 ± 2 | 114 | 3 | 35 ± 2 | 85 | 3 |
| Day 4 | 34 ± 2 | 109 | 3 | 32 ± 1 | 116 | 3 |
| Day 6 | 35 ± 1 | 109 | 3 | 34 ± 2 | 99 | 3 |
| $I_{mean}$ (AHA) (a.u.) | IL-1β | | | | | |
| | WT | | | *Dcn*$^{-/-}$ | | |
| | mean ± 95% CI | *n* (cells) | *N* (animals) | mean ± 95% CI | *n* (cells) | *N* (animals) |
| Day 2 | 37 ± 2 | 119 | 3 | 35 ± 2 | 119 | 3 |
| Day 4 | 37 ± 1 | 125 | 3 | 34 ± 2 | 113 | 3 |
| Day 6 | 33 ± 1 | 123 | 3 | 38 ± 2 | 123 | 3 |

| *p*-value (genotype/IL-1β) | WT vs *Dcn*$^{-/-}$ | | Untreated vs IL-1β | |
|---|---|---|---|---|
| | Untreated | IL-1β | WT | *Dcn*$^{-/-}$ |
| Day 0 | 0.762 | -- | -- | -- |
| Day 2 | 1.000 | 0.765 | 1.000 | 0.934 |
| Day 4 | 0.884 | 1.000 | 1.000 | 1.000 |
| Day 6 | 0.968 | 0.945 | 0.596 | 1.000 |

| *p*-value (time points) | Untreated | | IL-1β | |
|---|---|---|---|---|
| | WT | *Dcn*$^{-/-}$ | WT | *Dcn*$^{-/-}$ |
| Day 0 vs 2 | 0.995 | 0.997 | -- | -- |
| Day 0 vs 4 | 0.991 | 0.889 | -- | -- |
| Day 0 vs 6 | 0.993 | 1.000 | -- | -- |
| Day 2 vs 4 | 1.000 | 0.852 | 0.952 | 0.959 |
| Day 2 vs 6 | 0.948 | 1.000 | 0.702 | 0.906 |
| Day 4 vs 6 | 0.928 | 0.896 | 0.862 | 0.770 |

**Table S3.** Summary of averaged values and statistical analysis outcomes of the peak GAL intensity, $I_{peak}$, extracted from high-resolution confocal images of click-labeled femoral head cartilage explants of age-matched wild-type (WT) and decorin-null (*Dcn*$^{-/-}$) mice over 6 days of culture, shown as mean ± 95% CI from values averaged by each explant. The *p*-values were calculated using the linear mixed-effect model, followed by Holm-Bonferroni correction for multiple contrasts between genotypes and treatments, and Tukey-Kramer correction for multiple comparisons across culture time points for each genotype and treatment.

| $I_{peak}$ (GAL) (a.u.) | Untreated | | | | | |
|---|---|---|---|---|---|---|
| | WT | | | *Dcn*$^{-/-}$ | | |
| | mean ± 95% CI | *n* (cells) | *N* (animals) | mean ± 95% CI | *n* (cells) | *N* (animals) |
| Day 0 | 59 ± 2 | 114 | 4 | 56 ± 2 | 95 | 4 |
| Day 2 | 62 ± 2 | 78 | 4 | 48 ± 2 | 110 | 4 |
| Day 4 | 56 ± 2 | 111 | 4 | 46 ± 2 | 108 | 4 |
| Day 6 | 49 ± 2 | 104 | 4 | 37 ± 2 | 102 | 4 |
| $I_{peak}$ (GAL) (a.u.) | IL-1β | | | | | |
| | WT | | | *Dcn*$^{-/-}$ | | |
| | mean ± 95% CI | *n* (cells) | *N* (animals) | mean ± 95% CI | *n* (cells) | *N* (animals) |
| Day 2 | 45 ± 2 | 131 | 4 | 45 ± 2 | 111 | 4 |
| Day 4 | 33 ± 2 | 114 | 4 | 32 ± 2 | 109 | 4 |
| Day 6 | 25 ± 1 | 91 | 4 | 26 ± 1 | 97 | 4 |

| *p*-value (genotype/IL-1β) | WT vs *Dcn*$^{-/-}$ | | Untreated vs IL-1β | |
|---|---|---|---|---|
| | Control | IL-1β | WT | *Dcn*$^{-/-}$ |
| Day 0 | 0.307 | -- | -- | -- |
| Day 2 | 0.010 | 0.901 | 0.008 | 0.558 |
| Day 4 | 0.003 | 0.851 | < 0.001 | 0.001 |
| Day 6 | 0.012 | 0.865 | 0.001 | 0.001 |

| *p*-value (time points) | Untreated | | IL-1β | |
|---|---|---|---|---|
| | WT | *Dcn*$^{-/-}$ | WT | *Dcn*$^{-/-}$ |
| Day 0 vs 2 | 0.796 | 0.040 | -- | -- |
| Day 0 vs 4 | 0.521 | 0.008 | -- | -- |
| Day 0 vs 6 | 0.005 | < 0.001 | -- | -- |
| Day 2 vs 4 | 0.155 | 0.785 | 0.017 | 0.003 |
| Day 2 vs 6 | 0.001 | 0.008 | 0.001 | < 0.001 |
| Day 4 vs 6 | 0.050 | 0.040 | 0.187 | 0.158 |

**Table S4.** Summary of averaged values and statistical analysis outcomes of the matrix-to-cell Euclidean distance corresponding to the peak GAL intensity, $d_{peak}$, extracted from high-resolution confocal images of click-labeled femoral head cartilage explants of age-matched wild-type (WT) and decorin-null (*Dcn*$^{-/-}$) mice over 6 days of culture, shown as mean ± 95% CI from values averaged by each explant. The *p*-values were calculated using the linear mixed-effect model, followed by Holm-Bonferroni correction for multiple contrasts between genotypes and treatments, and Tukey-Kramer correction for multiple comparisons across culture time points for each genotype and treatment.

| $d_{peak}$ (GAL) (μm) | Untreated | | | | | |
|---|---|---|---|---|---|---|
| | WT | | | *Dcn*$^{-/-}$ | | |
| | mean ± 95% CI | *n* (cells) | *N* (animals) | mean ± 95% CI | *n* (cells) | *N* (animals) |
| Day 0 | 0.91 ± 0.06 | 114 | 4 | 0.92 ± 0.07 | 95 | 4 |
| Day 2 | 1.01 ± 0.09 | 78 | 4 | 1.03 ± 0.07 | 110 | 4 |
| Day 4 | 1.10 ± 0.08 | 111 | 4 | 1.07 ± 0.08 | 108 | 4 |
| Day 6 | 1.03 ± 0.07 | 104 | 4 | 1.05 ± 0.08 | 102 | 4 |
| $d_{peak}$ (GAL) (μm) | IL-1β | | | | | |
| | WT | | | *Dcn*$^{-/-}$ | | |
| | mean ± 95% CI | *n* (cells) | *N* (animals) | mean ± 95% CI | *n* (cells) | *N* (animals) |
| Day 2 | 0.92 ± 0.05 | 131 | 4 | 0.88 ± 0.07 | 111 | 4 |
| Day 4 | 0.91 ± 0.08 | 114 | 4 | 0.90 ± 0.06 | 109 | 4 |
| Day 6 | 1.00 ± 0.09 | 91 | 4 | 0.98 ± 0.09 | 97 | 4 |

| *p*-value (genotype/IL-1β) | WT vs *Dcn*$^{-/-}$ | | Untreated vs IL-1β | |
|---|---|---|---|---|
| | Control | IL-1β | WT | *Dcn*$^{-/-}$ |
| Day 0 | 0.983 | -- | -- | -- |
| Day 2 | 1.000 | 1.000 | 0.567 | 0.567 |
| Day 4 | 1.000 | 1.000 | 0.334 | 0.334 |
| Day 6 | 1.000 | 1.000 | 1.000 | 1.000 |

| *p*-value (time points) | Untreated | | IL-1β | |
|---|---|---|---|---|
| | WT | *Dcn*$^{-/-}$ | WT | *Dcn*$^{-/-}$ |
| Day 0 vs 2 | 0.837 | 0.665 | -- | -- |
| Day 0 vs 4 | 0.491 | 0.496 | -- | -- |
| Day 0 vs 6 | 0.873 | 0.457 | -- | -- |
| Day 2 vs 4 | 0.927 | 0.991 | 0.974 | 0.989 |
| Day 2 vs 6 | 1.000 | 0.982 | 0.600 | 0.797 |
| Day 4 vs 6 | 0.897 | 1.000 | 0.482 | 0.717 |

**Table S5.** Summary of averaged values and statistical analysis outcomes of the Euclidean distance from $d_{peak}$ to where the GAL intensity decays by 50%, $d_{1/2}$, extracted from high-resolution confocal images of click-labeled femoral head cartilage explants of age-matched wild-type (WT) and decorin-null (*Dcn*$^{-/-}$) mice over 6 days of culture, shown as mean ± 95% CI from values averaged by each explant. The *p*-values were calculated using the linear mixed-effect model, followed by Holm-Bonferroni correction for multiple contrasts between genotypes and treatments, and Tukey-Kramer correction for multiple comparisons across culture time points for each genotype and treatment.

| $d_{1/2}$ (GAL) (μm) | Untreated | | | | | |
|---|---|---|---|---|---|---|
| | WT | | | *Dcn*$^{-/-}$ | | |
| | mean ± 95% CI | *n* (cells) | *N* (animals) | mean ± 95% CI | *n* (cells) | *N* (animals) |
| Day 0 | 0.92 ± 0.08 | 114 | 4 | 0.82 ± 0.10 | 95 | 4 |
| Day 2 | 0.74 ± 0.11 | 78 | 4 | 0.78 ± 0.09 | 110 | 4 |
| Day 4 | 0.74 ± 0.09 | 111 | 4 | 0.72 ± 0.10 | 108 | 4 |
| Day 6 | 0.91 ± 0.09 | 104 | 4 | 0.76 ± 0.11 | 102 | 4 |
| $d_{1/2}$ (GAL) (μm) | IL-1β | | | | | |
| | WT | | | *Dcn*$^{-/-}$ | | |
| | mean ± 95% CI | *n* (cells) | *N* (animals) | mean ± 95% CI | *n* (cells) | *N* (animals) |
| Day 2 | 0.90 ± 0.07 | 131 | 4 | 0.85 ± 0.08 | 111 | 4 |
| Day 4 | 1.10 ± 0.13 | 114 | 4 | 0.99 ± 0.12 | 109 | 4 |
| Day 6 | 0.76 ± 0.13 | 91 | 4 | 1.15 ± 0.17 | 97 | 4 |

| *p*-value (genotype/IL-1β) | WT vs *Dcn*$^{-/-}$ | | Untreated vs IL-1β | |
|---|---|---|---|---|
| | Control | IL-1β | WT | *Dcn*$^{-/-}$ |
| Day 0 | 0.762 | -- | -- | ~ |
| Day 2 | 1.000 | 1.000 | 0.263 | 0.592 |
| Day 4 | 0.887 | 0.486 | 0.087 | 0.087 |
| Day 6 | 0.750 | 0.750 | 0.994 | 0.188 |

| *p*-value (time points) | Untreated | | IL-1β | |
|---|---|---|---|---|
| | WT | *Dcn*$^{-/-}$ | WT | *Dcn*$^{-/-}$ |
| Day 0 vs 2 | 0.698 | 0.976 | -- | -- |
| Day 0 vs 4 | 0.667 | 0.868 | -- | -- |
| Day 0 vs 6 | 0.996 | 0.988 | -- | -- |
| Day 2 vs 4 | 1.000 | 0.984 | 0.474 | 0.308 |
| Day 2 vs 6 | 0.820 | 1.000 | 0.922 | 0.032 |
| Day 4 vs 6 | 0.794 | 0.969 | 0.306 | 0.264 |

**Table S6.** Summary of averaged values and statistical analysis outcomes of the Euclidean distance from the cell surface to where the GAL intensity decays by 50%, $d_{peak} + d_{1/2}$, extracted from high-resolution confocal images of click-labeled femoral head cartilage explants of age-matched wild-type (WT) and decorin-null (*Dcn*$^{-/-}$) mice over 6 days of culture, shown as mean ± 95% CI from values averaged by each explant. The *p*-values were calculated using the linear mixed-effect model, followed by Holm-Bonferroni correction for multiple contrasts between genotypes and treatments, and Tukey-Kramer correction for multiple comparisons across culture time points for each genotype and treatment.

| $d_{peak} + d_{1/2}$ (GAL) (μm) | Untreated | | | | | |
|---|---|---|---|---|---|---|
| | WT | | | *Dcn*$^{-/-}$ | | |
| | mean ± 95% CI | *n* (cells) | *N* (animals) | mean ± 95% CI | *n* (cells) | *N* (animals) |
| Day 0 | 1.84 ± 0.09 | 114 | 4 | 1.73 ± 0.11 | 95 | 4 |
| Day 2 | 1.75 ± 0.13 | 78 | 4 | 1.81 ± 0.10 | 110 | 4 |
| Day 4 | 1.83 ± 0.09 | 111 | 4 | 1.79 ± 0.10 | 108 | 4 |
| Day 6 | 1.94 ± 0.11 | 104 | 4 | 1.82 ± 0.13 | 102 | 4 |
| $d_{peak} + d_{1/2}$ (GAL) (μm) | IL-1β | | | | | |
| | WT | | | *Dcn*$^{-/-}$ | | |
| | mean ± 95% CI | *n* (cells) | *N* (animals) | mean ± 95% CI | *n* (cells) | *N* (animals) |
| Day 2 | 1.82 ± 0.08 | 131 | 4 | 1.73 ± 0.10 | 111 | 4 |
| Day 4 | 1.99 ± 0.15 | 114 | 4 | 1.89 ± 0.13 | 109 | 4 |
| Day 6 | 1.76 ± 0.17 | 91 | 4 | 2.13 ± 0.18 | 97 | 4 |

| *p*-value (genotype/IL-1β) | WT vs *Dcn*$^{-/-}$ | | Untreated vs IL-1β | |
|---|---|---|---|---|
| | Control | IL-1β | WT | *Dcn*$^{-/-}$ |
| Day 0 | 0.726 | -- | -- | -- |
| Day 2 | 1.000 | 1.000 | 1.000 | 1.000 |
| Day 4 | 0.799 | 0.799 | 0.369 | 0.369 |
| Day 6 | 1.000 | 1.000 | 0.801 | 0.296 |

| *p*-value (time points) | Untreated | | IL-1β | |
|---|---|---|---|---|
| | WT | *Dcn*$^{-/-}$ | WT | *Dcn*$^{-/-}$ |
| Day 0 vs 2 | 0.983 | 0.965 | -- | -- |
| Day 0 vs 4 | 0.992 | 0.991 | -- | -- |
| Day 0 vs 6 | 0.902 | 0.853 | -- | -- |
| Day 2 vs 4 | 0.918 | 0.998 | 0.847 | 0.694 |
| Day 2 vs 6 | 0.742 | 0.986 | 0.951 | 0.121 |
| Day 4 vs 6 | 0.977 | 0.953 | 0.967 | 0.375 |

**Table S7.** Summary of averaged values and statistical analysis outcomes of the cumulative relative percentage of GAL released from click-labeled femoral head cartilage explants of age-matched wild-type (WT) and decorin-null (*Dcn*$^{-/-}$) mice over 6 days of culture under IL-1β stimulation, shown as mean ± 95% CI from values averaged by each explant. The *p*-values were calculated using the linear mixed-effect model, followed by Holm-Bonferroni correction for multiple contrasts between genotypes and treatments, and Tukey-Kramer correction for multiple comparisons across culture time points for each genotype and treatment.

| GAL release (%) | Untreated | | | |
|---|---|---|---|---|
| | WT | | *Dcn*$^{-/-}$ | |
| | mean ± 95% CI | *N* | mean ± 95% CI | *N* |
| Day 2 | 2.6 ± 0.8 | 6 | 4.5 ± 0.6 | 6 |
| Day 4 | 4.0 ± 0.9 | 6 | 6.3 ± 0.9 | 6 |
| Day 6 | 5.1 ± 1.0 | 6 | 7.6 ± 1.2 | 6 |
| **GAL release (%)** | **IL-1β** | | | |
| | WT | | *Dcn*$^{-/-}$ | |
| | mean ± 95% CI | *N* | mean ± 95% CI | *N* |
| Day 2 | 5.7 ± 0.4 | 6 | 6.8 ± 0.7 | 6 |
| Day 4 | 10.7 ± 0.8 | 6 | 13.0 ± 1.6 | 6 |
| Day 6 | 14.1 ± 1.1 | 6 | 17.2 ± 1.9 | 6 |

| *p*-value (genotype/IL-1β) | WT vs *Dcn*$^{-/-}$ | | Untreated vs IL-1β | |
|---|---|---|---|---|
| | Control | IL-1β | WT | *Dcn*$^{-/-}$ |
| Day 2 | 0.041 | 0.248 | < 0.001 | 0.035 |
| Day 4 | 0.014 | 0.028 | < 0.001 | < 0.001 |
| Day 6 | 0.009 | 0.009 | < 0.001 | < 0.001 |

| *p*-value (time points) | Untreated | | IL-1β | |
|---|---|---|---|---|
| | WT | *Dcn*$^{-/-}$ | WT | *Dcn*$^{-/-}$ |
| Day 2 vs Day 4 | < 0.001 | < 0.001 | < 0.001 | < 0.001 |
| Day 2 vs Day 6 | < 0.001 | < 0.001 | < 0.001 | < 0.001 |
| Day 4 vs Day 6 | < 0.001 | 0.002 | < 0.001 | < 0.001 |

**Table S8.** Summary of averaged values and statistical analysis outcomes of the cumulative relative percentage of AHA released from click-labeled femoral head cartilage explants of age-matched wild-type (WT) and decorin-null (*Dcn*$^{-/-}$) mice over 6 days under IL-1β stimulation, shown as mean ± 95% confidence intervals from values averaged by each explant. The *p*-values were calculated using the linear mixed-effect model, followed by Holm-Bonferroni correction for multiple contrasts between genotypes and treatments, and Tukey-Kramer correction for multiple comparisons across culture time points for each genotype and treatment.

| | Untreated | | | |
|---|---|---|---|---|
| | WT | | *Dcn*$^{-/-}$ | |
| | Mean ± 95% CI | N | Mean ± 95% CI | N |
| Day 2 | 5.1 ± 0.9 | 5 | 4.8 ± 0.5 | 6 |
| Day 4 | 7.1 ± 0.9 | 5 | 7.1 ± 0.5 | 6 |
| Day 6 | 8.4 ± 1.0 | 5 | 8.5 ± 0.6 | 6 |
| | IL-1β | | | |
| | WT | | *Dcn*$^{-/-}$ | |
| | Mean ± 95% CI | N | Mean ± 95% CI | N |
| Day 2 | 5.4 ± 1.8 | 5 | 4.9 ± 0.7 | 6 |
| Day 4 | 7.7 ± 1.9 | 5 | 7.7 ± 1.0 | 6 |
| Day 6 | 9.2 ± 2.2 | 5 | 9.4 ± 1.1 | 6 |

| *p*-value (genotype/IL-1β) | WT vs *Dcn*$^{-/-}$ | | Untreated vs IL-1β | |
|---|---|---|---|---|
| | Control | IL-1β | WT | *Dcn*$^{-/-}$ |
| Day 2 | 1.000 | 1.000 | 1.000 | 1.000 |
| Day 4 | 1.000 | 1.000 | 0.679 | 0.679 |
| Day 6 | 1.000 | 1.000 | 0.545 | 0.332 |

| *p*-value (time points) | Untreated | | IL-1β | |
|---|---|---|---|---|
| | WT | *Dcn*$^{-/-}$ | WT | *Dcn*$^{-/-}$ |
| Day 2 vs Day 4 | < 0.001 | < 0.001 | < 0.001 | < 0.001 |
| Day 2 vs Day 6 | < 0.001 | < 0.001 | < 0.001 | < 0.001 |
| Day 4 vs Day 6 | 0.007 | < 0.001 | < 0.001 | < 0.001 |

**Table S9.** Summary of averaged values and statistical analysis outcomes of the cumulative relative percentage of GAL released from click-labeled femoral head cartilage explants of age-matched wild-type (WT) and decorin knockout (*Dcn*$^{-/-}$) mice over 8 days of culture under IL-1β stimulation with the filtration of exogenous decorin either before or after the onset of IL-1β (Pre-Dcn, Post-Dcn, Comb-Dcn), shown as mean ± 95% CI from values averaged by each explant. The *p*-values were calculated using the linear mixed-effect model, followed by Holm-Bonferroni correction for multiple contrasts between genotypes, and Tukey-Kramer correction for multiple comparisons among different treatment conditions for each genotype and time point, and across culture time points for each genotype and treatment.

| | WT | | | | | | | | | |
|---|---|---|---|---|---|---|---|---|---|---|
| | Untreated | | IL-1β only | | Pre-Dcn | | Post-Dcn | | Comb-Dcn | |
| | mean ± 95% CI | *N* | mean ± 95% CI | *N* | mean ± 95% CI | *N* | mean ± 95% CI | *N* | mean ± 95% CI | *N* |
| Day 2 | 2.8 ± 0.6 | 7 | 8.1 ± 1.2 | 6 | 7.9 ± 1.2 | 10 | 7.9 ± 1.3 | 10 | 7.7 ± 1.1 | 5 |
| Day 4 | 5.1 ± 0.4 | 7 | 12.6 ± 0.7 | 6 | 12.2 ± 1.4 | 10 | 14.6 ± 1.3 | 10 | 13.2 ± 1.1 | 5 |
| Day 6 | 6.6 ± 0.3 | 7 | 16.6 ± 0.9 | 6 | 14.5 ± 1.2 | 10 | 19.2 ± 1.4 | 10 | 15.6 ± 1.1 | 5 |
| Day 8 | 8.1 ± 0.3 | 7 | 24.5 ± 2.2 | 6 | 16.4 ± 1.3 | 10 | 26.9 ± 2.3 | 10 | 17.6 ± 1.1 | 5 |
| | *Dcn*$^{-/-}$ | | | | | | | | | |
| | Untreated | | IL-1β only | | Pre-Dcn | | Post-Dcn | | Comb-Dcn | |
| | mean ± 95% CI | *N* | mean ± 95% CI | *N* | mean ± 95% CI | *N* | mean ± 95% CI | *N* | mean ± 95% CI | *N* |
| Day 2 | 3.9 ± 0.5 | 6 | 10.8 ± 1.8 | 7 | 9.4 ± 1.1 | 11 | 10.6 ± 0.9 | 8 | 9.8 ± 1.8 | 4 |
| Day 4 | 7.9 ± 1.1 | 6 | 15.7 ± 0.9 | 7 | 15.0 ± 1.3 | 11 | 16.4 ± 1.0 | 8 | 14.9 ± 1.8 | 4 |
| Day 6 | 9.7 ± 1.3 | 6 | 19.7 ± 1.8 | 7 | 20.3 ± 2.7 | 11 | 19.7 ± 1.3 | 8 | 18.5 ± 1.8 | 4 |
| Day 8 | 11.2 ± 1.7 | 6 | 28.4 ± 1.2 | 7 | 29.7 ± 1.3 | 11 | 31.2 ± 1.5 | 8 | 32.0 ± 2.4 | 4 |

| *p*-value (treatment conditions) | | WT | | | | *Dcn*$^{-/-}$ | | | |
|---|---|---|---|---|---|---|---|---|---|
| | | IL-1β only | Pre-Dcn | Post-Dcn | Comb-Dcn | IL-1β only | Pre-Dcn | Post-Dcn | Comb-Dcn |
| Day 2 | Untreated | < 0.001 | < 0.001 | < 0.001 | < 0.001 | < 0.001 | < 0.001 | < 0.001 | 0.002 |
| | IL-1β only | -- | 1.000 | 1.000 | 0.998 | -- | 1.000 | 0.720 | 0.946 |
| | Pre-Dcn | -- | -- | 1 | 1.000 | -- | -- | 0.808 | 0.974 |
| | Post-Dcn | -- | -- | -- | 1.000 | -- | -- | -- | 0.999 |
| Day 4 | Untreated | < 0.001 | < 0.001 | < 0.001 | < 0.001 | < 0.001 | < 0.001 | < 0.001 | < 0.001 |
| | IL-1β only | -- | 0.993 | 0.338 | 0.998 | -- | 0.976 | 0.971 | 0.984 |
| | Pre-Dcn | -- | -- | 0.071 | 0.884 | -- | -- | 0.688 | 0.836 |
| | Post-Dcn | -- | -- | -- | 0.735 | -- | -- | -- | 1 |
| Day 6 | Untreated | < 0.001 | < 0.001 | < 0.001 | < 0.001 | < 0.001 | < 0.001 | < 0.001 | < 0.001 |
| | IL-1β only | -- | 0.303 | 0.106 | 0.943 | -- | 1 | 0.983 | 0.935 |
| | Pre-Dcn | -- | -- | < 0.001 | 0.856 | -- | -- | 0.982 | 0.925 |
| | Post-Dcn | -- | -- | -- | 0.018 | -- | -- | -- | 0.699 |
| Day 8 | Untreated | < 0.001 | < 0.001 | < 0.001 | < 0.001 | < 0.001 | < 0.001 | < 0.001 | < 0.001 |
| | IL-1β only | -- | < 0.001 | 0.147 | < 0.001 | -- | 0.1491 | 0.765 | 0.111 |
| | Pre-Dcn | -- | -- | < 0.001 | 0.790 | -- | -- | 0.652 | 0.981 |
| | Post-Dcn | -- | -- | -- | < 0.001 | -- | -- | -- | 0.460 |

**Table S9.** (Continued)

| *p*-value (genotypes) | WT vs *Dcn*$^{-/-}$ | | | | |
|---|---|---|---|---|---|
| | Untreated | IL-1β only | Pre-Dcn | Post-Dcn | Comb-Dcn |
| Day 2 | 0.312 | 0.072 | 0.047 | 0.312 | 0.312 |
| Day 4 | 0.002 | 0.029 | 0.002 | 0.774 | 0.415 |
| Day 6 | < 0.001 | 0.024 | < 0.001 | 0.366 | 0.080 |
| Day 8 | < 0.001 | 0.002 | < 0.001 | 0.030 | < 0.001 |

| *p*-value (time points) | WT | | | | |
|---|---|---|---|---|---|
| | Untreated | IL-1β only | Pre-Dcn | Post-Dcn | Comb-Dcn |
| Day 2 vs 4 | < 0.001 | 0.002 | < 0.001 | < 0.001 | < 0.001 |
| Day 2 vs 6 | < 0.001 | < 0.001 | < 0.001 | < 0.001 | < 0.001 |
| Day 2 vs 8 | < 0.001 | < 0.001 | < 0.001 | < 0.001 | < 0.001 |
| Day 4 vs 6 | < 0.001 | 0.005 | < 0.001 | 0.003 | < 0.001 |
| Day 4 vs 8 | < 0.001 | < 0.001 | < 0.001 | < 0.001 | < 0.001 |
| Day 6 vs 8 | < 0.001 | < 0.001 | < 0.001 | < 0.001 | 0.004 |
| *p*-value (time points) | *Dcn*$^{-/-}$ | | | | |
| | Untreated | IL-1β only | Pre-Dcn | Post-Dcn | Comb-Dcn |
| Day 2 vs 4 | < 0.001 | 0.002 | < 0.001 | < 0.001 | < 0.001 |
| Day 2 vs 6 | < 0.001 | < 0.001 | < 0.001 | < 0.001 | < 0.001 |
| Day 2 vs 8 | < 0.001 | < 0.001 | < 0.001 | < 0.001 | < 0.001 |
| Day 4 vs 6 | 0.065 | 0.007 | < 0.001 | < 0.001 | 0.006 |
| Day 4 vs 8 | 0.001 | < 0.001 | < 0.001 | < 0.001 | < 0.001 |
| Day 6 vs 8 | 0.148 | < 0.001 | < 0.001 | < 0.001 | < 0.001 |

**Table S10.**: Summary of fitting outcomes for the ternary bindings measured by surface plasmon resonance (SPR), including the apparent association and dissociation rate constants ($k_a$ and $k_d$), as well as the equilibrium dissociation constant ($K_D$), shown for values of each individual replicate calculated from Scrubber, as well as mean ± SD for the fold changes of $1/K_D$ measured on each individual experiment with versus without decorin as the coupler repeated for $n$ = 3 replicates.

| Ligand | Analyte | Without Decorin | | | With Decorin | | | Fold change in $1/K_D$ | mean ± SD of fold change in $1/K_D$ |
|---|---|---|---|---|---|---|---|---|---|
| | | $k_a$ ($\times 10^4$ $M^{-1}\cdot s^{-1}$) | $k_d$ ($\times 10^{-3}$ $s^{-1}$) | $K_D$ (nM) | $k_a$ ($\times 10^4$ $M^{-1}\cdot s^{-1}$) | $k_d$ ($\times 10^{-3}$ $s^{-1}$) | $K_D$ (nM) | | |
| Aggrecan | Aggrecan | 27 ± 2 | (9.5 ± 0.1) × $10^2$ | (3.5 ± 0.3) × $10^3$ | 67 ± 3 | (1.1 ± 0.1) × $10^3$ | (1.6 ± 0.1) × $10^3$ | 2.20 | 2.7 ± 0.8 |
| | | (8.8 ± 0.5) × $10^2$ | (6.0 ± 0.1) × $10^2$ | 68 ± 1 | (2.1 ± 0.1) × $10^3$ | (6.7 ± 0.1) × $10^2$ | 31 ± 1 | 2.19 | |
| | | (2.9 ± 0.1) × $10^2$ | (4.4 ± 0.1) × $10^2$ | (1.5 ± 0.1) × $10^2$ | (7.5 ± 0.6) × $10^2$ | (3.1 ± 0.1) × $10^2$ | 42 ± 2 | 3.62 | |
| Collagen | Aggrecan | 4.1 ± 0.1 | 15.4 ± 0.2 | (3.7 ± 0.1) × $10^2$ | 5.0 ± 0.1 | 7.9 ± 0.1 | 157 ± 4 | 2.36 | 2.3 ± 0.1 |
| | | (1.7 ± 0.1) × $10^3$ | (6.4 ± 0.1) × $10^2$ | 36 ± 1 | (4.4 ± 0.1) × $10^3$ | (6.5 ± 0.1) × $10^2$ | 14.7 ± 0.2 | 2.46 | |
| | | 8.9 ± 0.1 | 2.7 ± 0.1 | 30 ± 1 | 10.0 ± 0.1 | 1.38 ± 0.03 | 13.7 ± 0.2 | 2.20 | |
| Aggrecan | Collagen | 8.2 ± 0.3 | (5.3 ± 0.1) × $10^2$ | (6.5 ± 0.2) × $10^3$ | 23 ± 1 | (4.1 ± 0.1) × $10^2$ | (1.8 ± 0.1) × $10^3$ | 3.57 | 3.4 ± 1.6 |
| | | 15.1 ± 0.8 | (6.1 ± 0.1) × $10^2$ | (4.0 ± 0.2) × $10^3$ | (1.1 ± 0.1) × $10^2$ | (8.6 ± 0.1) × $10^2$ | (8.2 ± 0.2) × $10^2$ | 4.88 | |
| | | 1.4 ± 0.2 | (6.3 ± 0.1) × $10^2$ | (4.6 ± 0.6) × $10^4$ | 2.2 ± 0.2 | (6.1 ± 0.1) × $10^2$ | (2.7 ± 0.1) × $10^4$ | 1.70 | |
| Collagen | Collagen | 24 ± 1 | 0.8 ± 0.1 | 3.4 ± 0.5 | 30 ± 1 | 0.43 ± 0.01 | 1.4 ± 0.1 | 2.33 | 1.7 ± 0.6 |
| | | 27 ± 1 | 4.6 ± 0.1 | 17 ± 1 | 26 ± 4 | 3.5 ± 0.1 | 13.5 ± 0.2 | 1.27 | |
| | | 35 ± 1 | 8.3 ± 0.1 | 24 ± 1 | 29 ± 1 | 4.4 ± 0.1 | 15.1 ± 0.2 | 1.58 | |

**Table S11.** Summary of averaged values and statistical analysis outcomes of the indentation modulus, $E_{ind}$, of femoral condylar cartilage from age-matched wild-type (WT) and decorin-null (*Dcn*$^{-/-}$) mice subjected to enzymatic GAG-depletion and re-infiltration with exogenous decorin, shown as mean ± 95% CI from values averaged by animal. The *p*-values were calculated using the linear mixed-effect model, followed by Holm-Bonferroni correction for multiple contrasts between genotypes and IL-1β stimulations, and Tukey-Kramer correction for multiple comparisons across different digestion and decorin infiltration conditions.

| $E_{ind}$ (kPa) | | − IL-1β | | | | + IL-1β | | |
|---|---|---|---|---|---|---|---|---|
| | | Undigested | Undigested + Dcn | GAG-dep. | GAG-dep. + Dcn | Undigested | GAG-dep. | GAG-dep. + Dcn |
| WT | mean ± 95% CI | 397 ± 81 | 310 ± 77 | 122 ± 17 | 253 ± 65 | 160 ± 21 | 152 ± 50 | 133 ± 16 |
| | *N* | 12 | 8 | 17 | 8 | 9 | 5 | 5 |
| *Dcn*$^{-/-}$ | mean ± 95% CI | 159 ± 37 | 153 ± 20 | 142 ± 33 | 149 ± 30 | 158 ± 37 | 155 ± 43 | 150 ± 31 |
| | *N* | 12 | 6 | 11 | 14 | 7 | 8 | 9 |
| *p*-value (genotype) | | < 0.001 | 0.009 | 0.108 | 0.012 | 1.000 | 1.000 | 1.000 |

| *p*-value (treatments) | WT | | | |
|---|---|---|---|---|
| | Undigested vs GAG-dep. | Undigested vs Undigested + Dcn | Undigested vs GAG-dep. + Dcn | GAG-dep. vs GAG-dep. + Dcn |
| − IL-1β | < 0.001 | 0.524 | 0.019 | 0.015 |
| + IL-1β | 0.897 | -- | 0.971 | 0.991 |
| *p*-value (treatments) | *Dcn*$^{-/-}$ | | | |
| | Undigested vs GAG-dep. | Undigested vs Undigested + Dcn | Undigested vs GAG-dep. + Dcn | GAG-dep. vs GAG-dep. + Dcn |
| − IL-1β | 0.751 | 0.751 | 0.751 | 0.751 |
| + IL-1β | 0.996 | -- | 0.960 | 0.981 |

| *p*-value (IL-1β) | Undigested | GAG-dep. | GAG-dep. + Dcn |
|---|---|---|---|
| WT | < 0.001 | 0.084 | 0.048 |
| *Dcn*$^{-/-}$ | 1.000 | 1.000 | 1.000 |

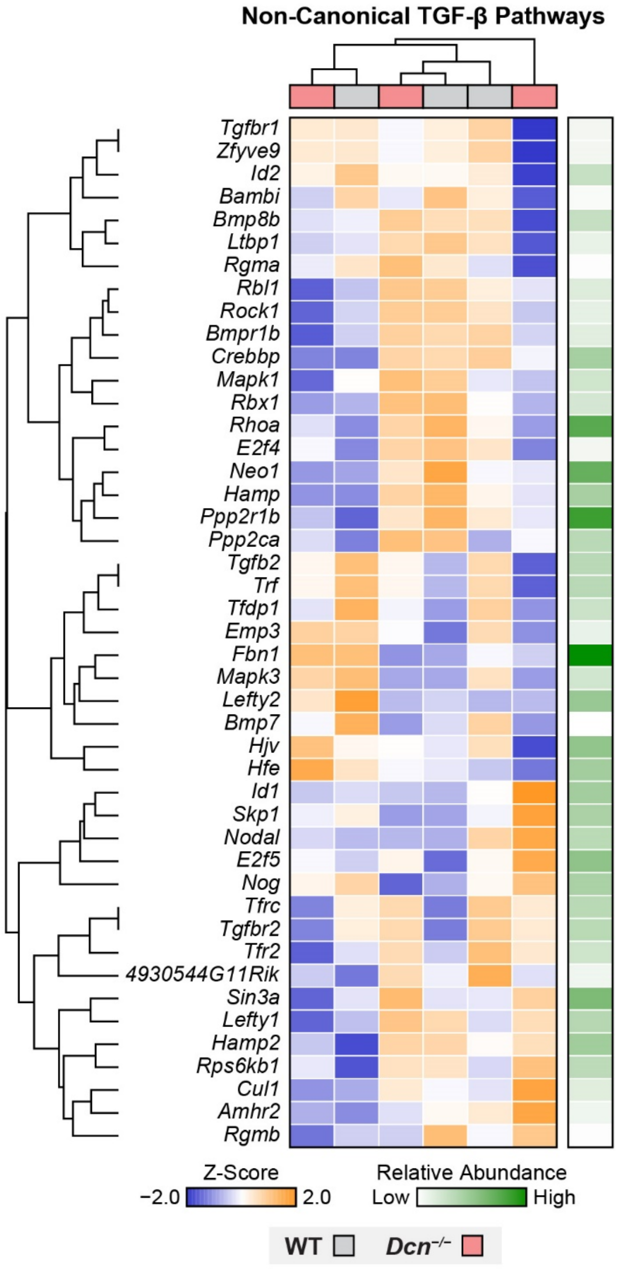


**Figure S1.** Heatmap of unbiased clustering for genes involved in KEGG non-canonical TGF-β signaling pathways show no clear separation in signaling activities between 3-month-old WT and *Dcn*$^{-/-}$ cartilage ($n$ = 3 per genotype).

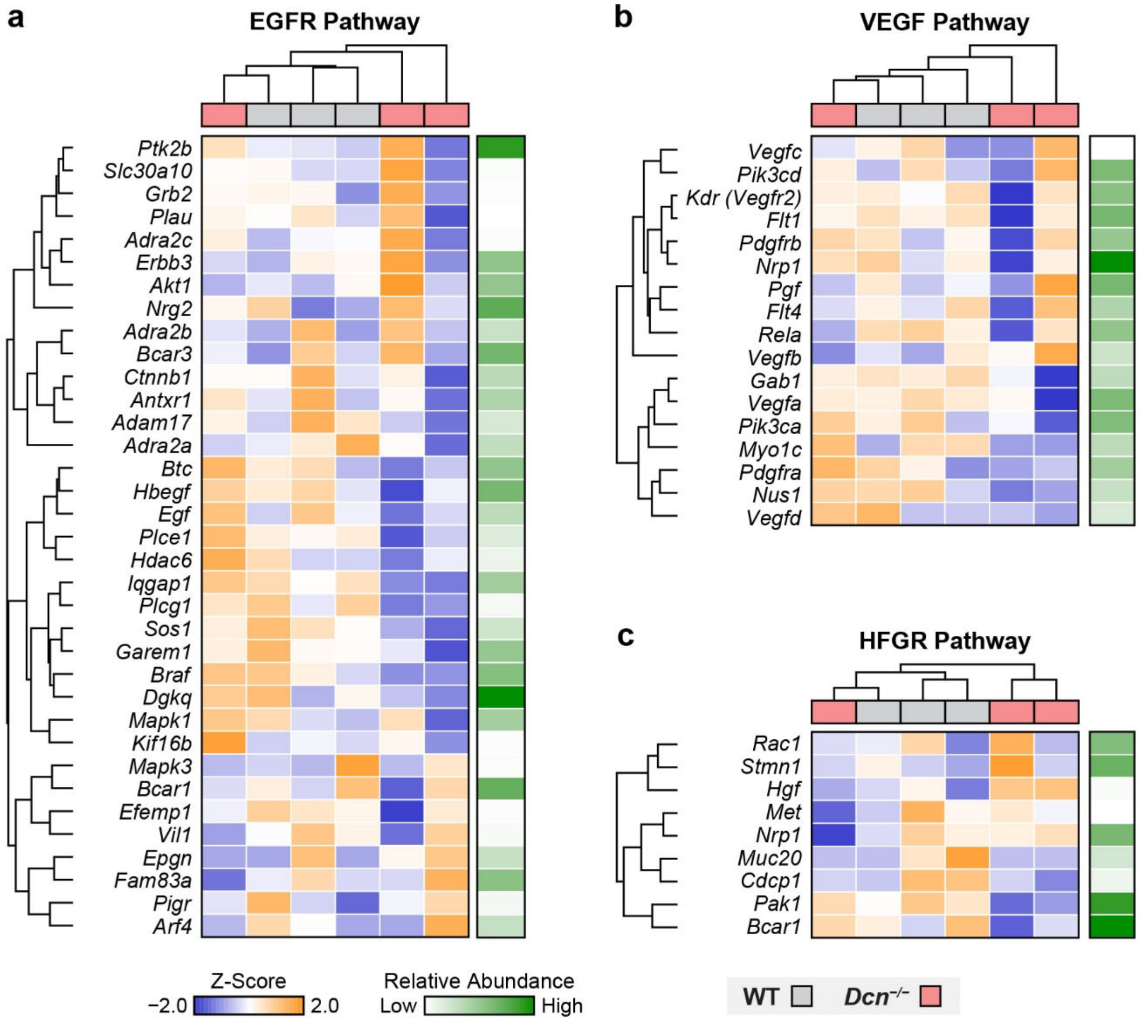


**Figure S2.** Heatmap of unbiased clustering for genes involved in other pathways potentially mediated by decorin from the GO database show no clear separation in these signaling activities between 3-month-old WT and *Dcn*$^{-/-}$ cartilage (*n* = 3 per genotype): a) EGFR pathway, b) VEGF pathway, c) HFGR pathway.

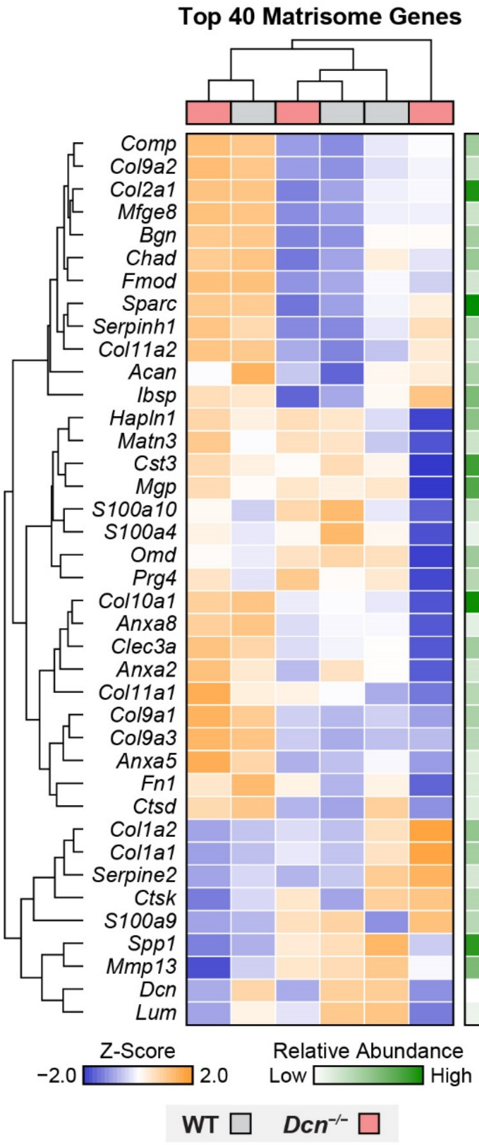


**Figure S3.** Heatmap of unbiased clustering of top 40 genes from the GO Matrisome Database reveals no clear separation in matrix biosynthesis activities between WT and *Dcn*$^{-/-}$ cartilage (*n* = 3 genotype).

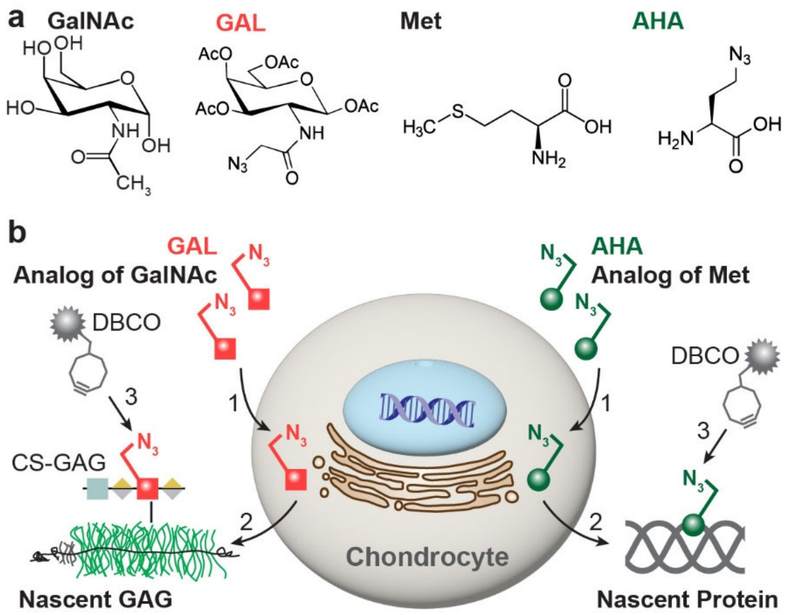


**Figure S4.** Schematic illustration of click-labeling for nascent glycosaminoglycans (GAGs) and proteins. a) Molecular structures of *N*-acetylgalactosamine (GalNAc), *N*-azidoacetylgalactosaminetetraacylated (GAL), methionine (Met), and azidohomoalanine (AHA). b) The azide-modified monosaccharide GAL serves as an analog of GalNAc and is metabolically incorporated into newly synthesized GAGs. Similarly, azidohomoalanine AHA, an analog of methionine, is incorporated into newly synthesized proteins. The click-labeling reagent, e.g., dibenzocyclooctyne (DBCO), is then applied to label GAL-containing nascent GAGs and AHA-containing nascent proteins via bio-orthogonal copper-free click chemistry, enabling their visualization and quantitative analysis.

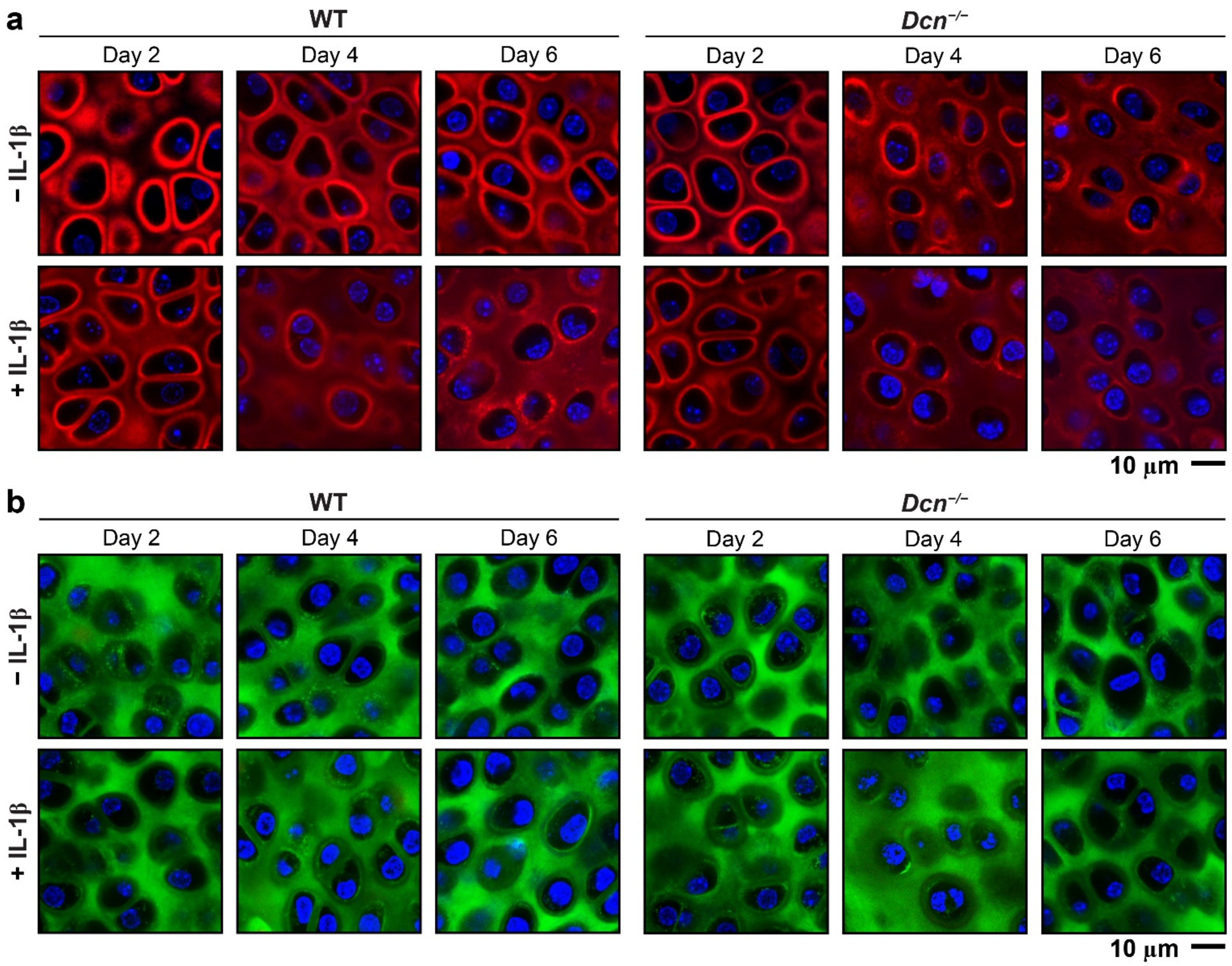


**Figure S5.** Representative confocal images of GAL (red) and AHA (green) labeling in 3-week-old wild-type (WT) and decorin-null (*Dcn*$^{-/-}$) cartilage explants cultured for 2, 4 and 6 days with or without IL-1β stimulation. Results show preferential localization of nascent GAGs in the pericellular matrix (PCM) and widespread distribution of nascent proteins throughout the extracellular matrix (ECM).

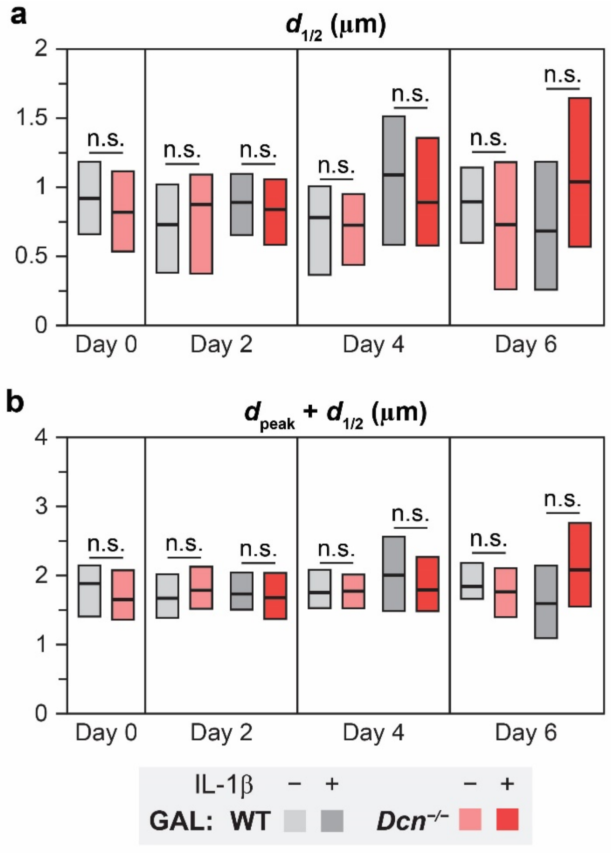


**Figure S6.** Loss of decorin does not affect additional spatial parameters of GAL signals between wild-type (WT) and decorin-null (*Dcn*$^{-/-}$) cartilage from day 0 to day 6 of culture with or without IL-1β: a) $d_{1/2}$ and b) $d_{peak} + d_{1/2}$. Box plots represent data obtained from > 75 ROIs across $n \geq 4$ animals for each group, n.s.: not significant.

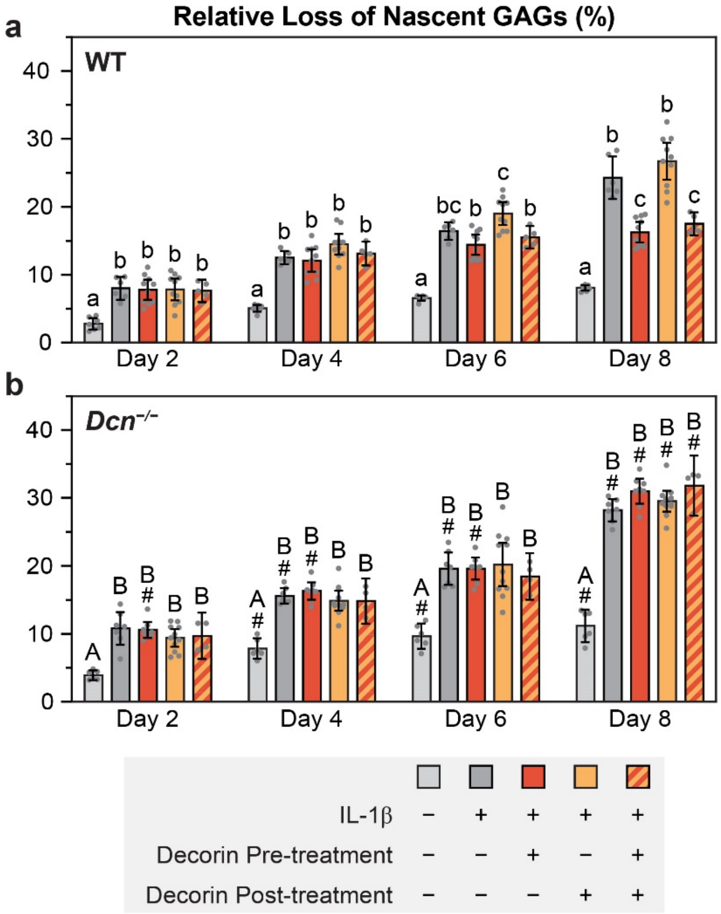


**Figure S7.** Cumulative release of nascent GAGs from a) WT and b) *Dcn*$^{-/-}$ cartilage explants at days 2, 4, 6 and 8 of culture under various decorin treatment conditions with IL-1β stimulation. Each data point represents one biological replicate (mean ± 95% CI, $n \geq 4$). Different letters indicate significant differences among treatments within each genotype and time point. $^{\#}$: $p < 0.05$ between genotypes from the same treatment and time point.
